# Phosphorylation of the Kv7.4 B helix reorganizes calmodulin interactions and reduces PIP_2_ binding

**DOI:** 10.64898/2026.09.18.752758

**Authors:** Phuc Phan, Courage Okose, Divya Sapkota, Willy Munyao, Dylan Girodat, Crystal R. Archer

**Author notes:** P.P., Department of Chemistry and Biochemistry, University of Delaware, Newark, DE; for W.M., ^3^Department of Translational Neuroscience, Barrow Neurological Institute, Phoenix, AR. To whom correspondence should be addressed: **Crystal R. Archer, PhD**: Department of Chemistry and Biochemistry, University of Arkansas, Fayetteville, AR.

## Abstract

Voltage-gated Kv7 (KCNQ, M-type) potassium channels regulate cellular excitability through interactions with phosphatidylinositol 4,5-bisphosphate (PIP_2_) and calmodulin (CaM). The distal B helix of Kv7.2-5 channels contains a conserved protein kinase C (PKC) phosphorylation site, suggesting that phosphorylation may regulate channel activity by altering CaM- and PIP_2_-dependent mechanisms. Here, we investigated the effects of phosphorylation of Thr552 within the Kv7.4 B helix using electrophysiology, biophysical assays, NMR spectroscopy, and molecular dynamics simulations. Phosphomimetic substitution of Thr552 produced a loss-of-function phenotype characterized by reduced current density and a depolarizing shift in voltage-dependent activation without altering channel surface expression. Biophysical studies demonstrated that phosphorylation produced only modest effects on overall CaM association. However, NMR spectroscopy revealed phosphorylation-dependent changes in apoCaM interactions, and molecular dynamics simulations identified remodeling of the local interaction network surrounding the distal B-helix polybasic “KRK motif” region. In contrast, phosphorylation markedly reduced PIP_2_ binding to both the isolated B helix and CaM-associated Kv7.4 regulatory complexes. Mutation of the distal B-helix KRK motif similarly impaired PIP_2_ binding, identifying this region as an important determinant of lipid recognition. Together, these findings support a model in which phosphorylation of Thr552 suppresses Kv7.4 activity primarily by reducing PIP_2_ interactions while reorganizing, rather than disrupting, CaM binding. These results identify Thr552 as a critical regulatory site linking phosphorylation-dependent signaling to Kv7.4 channel gating.

## INTRODUCTION

Voltage-gated Kv7(KCNQ) potassium channels comprise a family of five homologous subunits (Kv7.1-5) that assemble as homo- or heterotetrameric channels to regulate membrane excitability in numerous tissues^1^. Kv7.4 is highly expressed in cochlear outer hair cells, vascular and airway smooth muscle, neurons and epithelial tissues, where they regulate membrane excitability and potassium homeostasis^2,3^. Dysfunction of Kv7.4 channels causes progressive hearing loss and other disorders of excitability^4,5^. Channel activity depends on coordinated regulation by the membrane phospholipid phosphatidylinositol 4,5-bisphosphate (PIP_2_) and the Ca^2+^-sensor protein calmodulin (CaM)^6,7^. CaM associates with two conserved helices within the C terminus, the A and B helices, and contributes to channel assembly, trafficking and gating^6,8,9^. PIP_2_ interacts with multiple sites throughout Kv7 channels, including regions within the voltage-sensing domain, pore domain, and intracellular regulatory domain^7,10^. The distal B helix contains a conserved polybasic region that forms part of the CaM binding interface in crystal structures of the CaM-Kv7 complex and has been implicated in CaM-dependent channel regulation^11,12^. In Kv7.4, this region includes residues K546-R547-K548 (KRK motif) and lies immediately adjacent to Thr552. Because analogous distal B helix regions have been implicated in PIP2-dependent regulation in other Kv7 subunits^9–11,13,14^, phosphorylation at this site may influence local electrostatic interactions involved in channel regulation. Direct evidence for PIP_2_ binding to this region in Kv7.4, however, is limited. Whereas the A helix contains a canonical CaM-binding IQ-motif typically known to bind apoCaM, the B helix contains multiple proposed CaM binding motifs, including 1-10, 1-5-10, 1-12, 1-14 and 1-16 binding positions, some of which do not include the KRK motif or Thr552 **(Figure S1)**^14,15^. Multiple binding positions supports the hypothesis that the distal B helix region functions as a regulatory hub integrating CaM binding, PIP_2_ interactions and phosphorylation-dependent signaling. Defining how PIP_2_ and CaM interact with the Kv7.4 B helix is therefore important for understanding mechanisms that regulate Kv7.4 channel function in health and disease.

Signaling from G_q/11_ receptors suppresses Kv7 activity through depletion of PIP_2_, elevation of intracellular Ca^2+^ and activation of protein kinase C (PKC)^16,17^. Unlike Kv7.1, Kv7.2-5 subunits contain conserved PKC phosphorylation site within the distal B helix. In Kv7.4, this site corresponds to Thr552, which lies adjacent to the polybasic region proposed to participate in CaM and PIP_2_ interactions. Previous studies demonstrated that phosphorylation of this conserved residue in Kv7.2 channels reduces channel activity and increases sensitivity to PIP_2_ depletion^12,18,19^; however, the molecular consequences of Thr552 phosphorylation are poorly understood.

Recent cryo-EM structures revealed multiple PIP_2_ interacting regions with Kv7.4 channels and showed that Thr552 lies near the interface formed by CaM binding together the A and B helices **(Figures 1 and S2)**^20^. These observations led to the prevailing hypothesis that phosphorylation at this site may alter CaM association and disrupt channel regulation. Whether phosphorylation of Thr552 directly affects CaM binding, PIP_2_ binding or both, is unknown.

**Figure 1.**
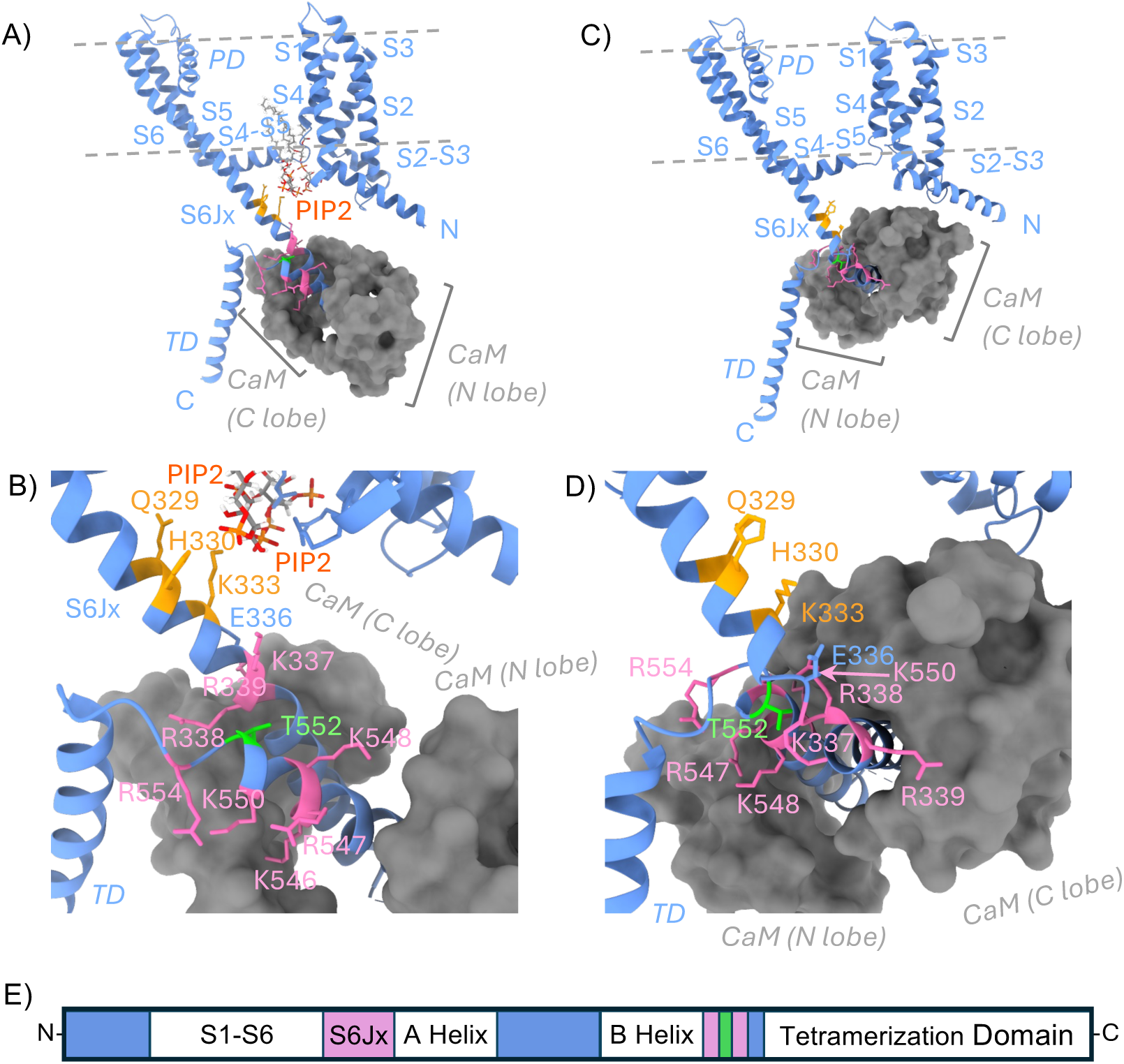
Sequence and orientation of the CaM-PIP_2_ binding domains of KCNQ4. Structural comparison of the PIP_2_-bound **(A, B)** and PIP_2_-free structure, (**C, D)** Kv7.4 complexes. A single Kv7.4 subunit is shown in blue and CaM in gray. Panels B and D provide enlarged highlighting the position and orientation of Thr552 within the CaM-PIP_2_ interface. Residues within the S6Jx region that interact with PIP_2_ are colored orange, cationic residues adjacent to the A and B helices colored pink, and Thr552 is colored green. **(E)** *S*chematic representation of the A and B helices within a linearized Kv7.4 subunit. Peptide sequences used in this study are shown below the schematic and are based on UniProt entry P56696-1. Residues highlighted in pink indicate PIP_2_ binding regions that extend beyond the CaM-binding domains, and Thr552 is highlighted in green.

Here we investigated how phosphorylation of Thr552 influences Kv7.4 regulation using electrophysiology, biochemical assays, NMR spectroscopy, molecular dynamics simulations, and quantitative binding measurements. We show that phosphorylation of Thr552 produces a strong loss-of-function phenotype without altering channel surface expression or substantially disrupting CaM binding affinity. Instead, phosphorylation remodels the local interaction network surrounding the distal B helix residue Lys548 and markedly reduces PIP_2_ interactions with the Kv7.4 regulatory domain. These findings identify a phosphorylation-dependent mechanism that preservers overall CaM association while altering the manner in which CaM and PIP_2_ interact with the Kv7.4 regulatory domain.

## RESULTS

### 2.1. Phosphorylation of Thr552 produces a calcium-independent loss-of-function phenotype

To determine the functional consequences of phosphorylation at Thr552 on Kv7.4 channel activity, we employed whole-cell patch-clamp electrophysiology using three channel variants transfected in HEK293T cells: Wild-type (WT), the phosphorylation-deficient T552A mutant (TA), and the phosphomimetic T552D mutant (TD). Channels were expressed either alone or together with wild-type calmodulin (CaM), and recordings were performed using intracellular solutions containing either EGTA (low Ca^2+^) or 50 μM Ca^2+^ to assess calcium-dependent regulation mediated by calmodulin. The activation parameters were consistent with previously reported values for Kv7.4 channels^21^. The data are summarized in **Figure 2** and supplemental table **ST1**.

**Figure 2.**
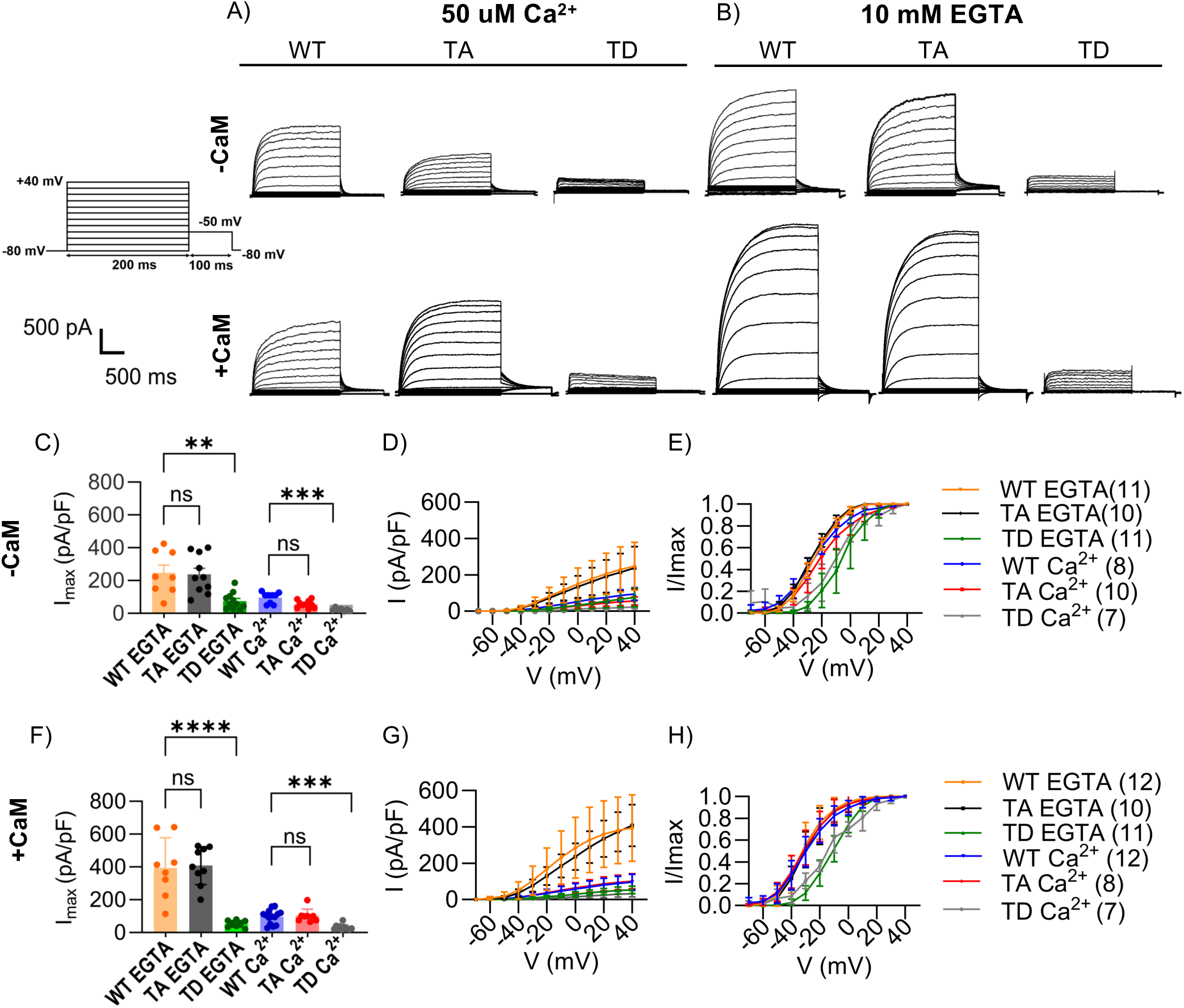
Phosphorylation at Thr552 reduces the activity of Kv7.4. **(A, B)** Representative whole-cell current traces from cells expressing Kv7.4-WT, T552A (TA), and T552D (TD) using pipette solutions containing either 50 µM Ca^2+^ (high intracellular Ca^2+^), or 10 mM EGTA (low intracellular Ca^2+^). **(C)** Summary of average current densities measured at +40 mV for Kv7.4-WT, T552A, and T552D under both EGTA and Ca^2+^ conditions in the absence of CaM overexpression. **(D)** Current voltage (I-V) relationships for for Kv7.4-WT, T552A, and T552D expressed in the absence of CaM overexpression. **(E)** Voltage-dependence of activation curves derived from tail current analysis showing a depolarizing shift in V_0.5 act_ of the T552D mutant in both Ca^2+^ and EGTA in the absence of CaM overexpression. **(F)** measured at +40 mV for Kv7.4-WT, T552A, and T552D under both EGTA and Ca^2+^ conditions with CaM overexpression. **(G)** Current voltage (I-V) relationships for for Kv7.4-WT, T552A, and T552D expressed with CaM overexpression. **(H)** Voltage-dependence of activation curves derived from tail current analysis showing a depolarizing shift in V_0.5 act_ of the T552D mutant in both Ca^2+^ and EGTA with CaM overexpression. Data are represented as mean ± SD. The number of cells analyzed is indicated in the corresponding panels. Statistics significance was computed using one-way ANOVA followed by Bonferroni’s multiple comparisons test.

Under elevated Ca^2+^, in Kv7.4-expressing cells without overexpressed CaM, the WT channels exhibited the largest current amplitudes, whereas TA exhibited intermediate current amplitudes. The TD mutant displayed a significantly reduced current amplitude, compared to the WT and TA. Quantification of current density measured at +40 mV (pA/pF) confirmed these observations (**Figure 2C**). When the channel and CaM were co-expressed under high Ca^2+^ conditions, the TA and WT variants exhibited current amplitudes similar to WT without CaM, and the TD mutant consistently showing markedly attenuated current amplitudes (**Figure 2A**, lower panel).

Under low Ca^2+^ conditions (10 mM EGTA), the current amplitudes of Kv7.4 WT and TA were enhanced (∼2-3x higher) with and without CaM overexpression, but the phosphomimetic still significantly reduced the current amplitudes for the Kv7.4 channels **(Figure 2B)**.

In all conditions, the phosphomimetic TD mutants induced a right shift of the the current voltage (I-V) relationship (pA/pF) and voltage-dependent of activation curves (I_max_) **(Figures 2D-E, G-H)**. These results suggest that the phosphomimetic mutation primarily alters the voltage dependence of Kv7.4 activation by shifting channel activation toward more depolarized membrane potential. Together, these data show that phosphorylation of Thr552 strongly reduces Kv7.4 channel activity that appears to be independent of CaM or Ca^2+^ levels.

### 2.2. Phosphorylation of Thr552 does not alter channel surface expression

To rule out whether the T552D phosphomimetic had an effect on the channel trafficking, we examined the surface expression of the Kv7-transfected cells in high and low Ca^2+^ in the presence and absence of CaM. In the first experiment, we utilized a surface-biotinylation assay for HEK293T cells transfected with CaM-CFP, GFP and myc-tagged Kv7.4-WT, TA and TD variants. Biotinylated surface proteins were isolated with NeutrAvidin slurry then immunoblotted with anti-myc to identify levels of myc-tagged Kv7.4 in three independent experiments. Samples were also probed with antibodies targeting endogenous membrane Na^+^/K^+^-ATPase, and GFP to validate the membrane elution (ELU) fraction was not contaminated with cytosolic proteins. (**Figure 3A**). Similar levels of membrane Kv7.4 was observed in all conditions, with quantification of the surface proteins showing no significance **(Figure 3B)**.

**Figure 3.**
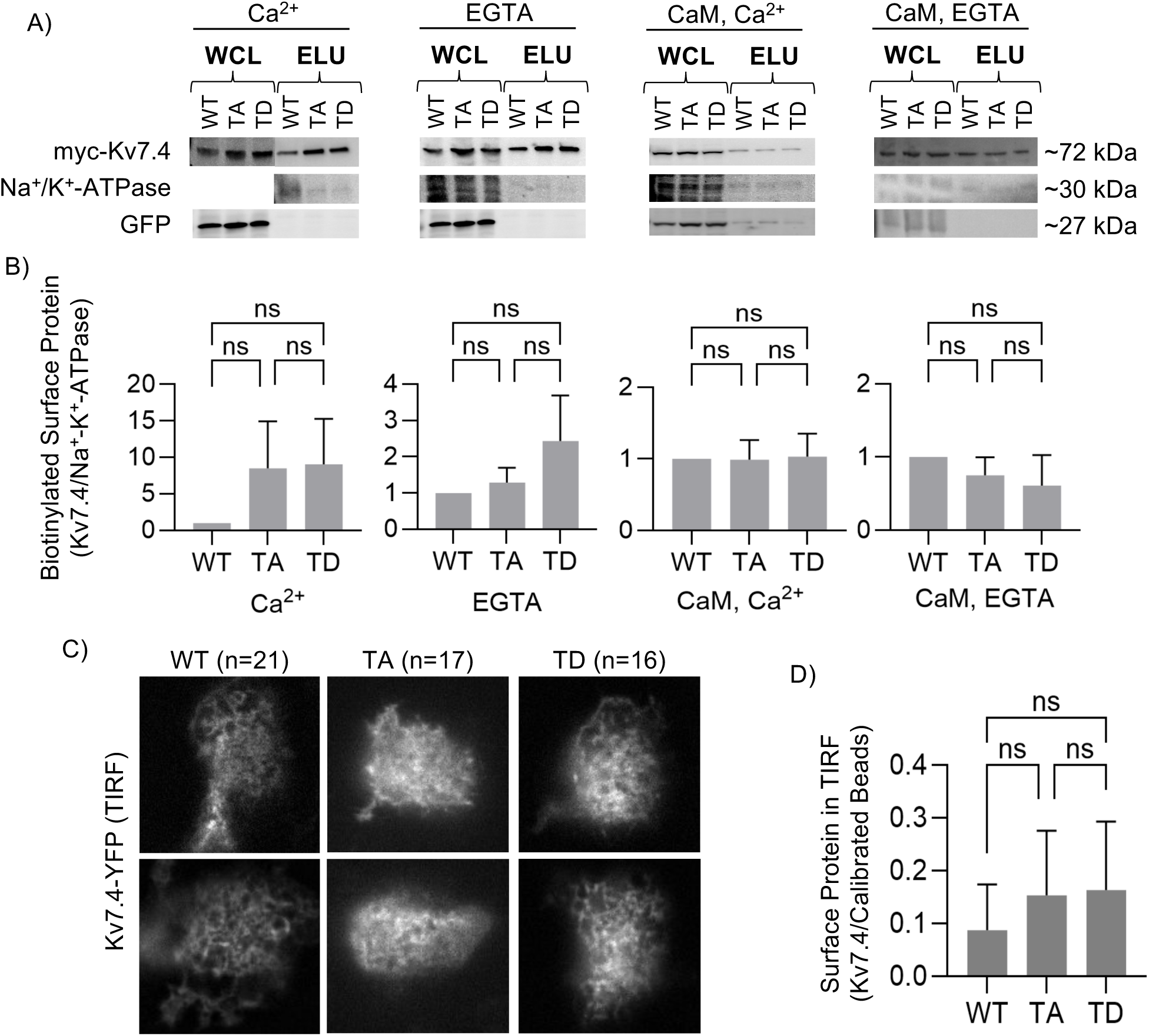
Surface Expression of Kv7.4 in HEK293T. **(A)** Representative immunoblots from surface biotinylation experiments (n=3 independent experiments) performed in HEK293T cells expressing Kv7.4-WT, T552A (TA) or T552D (TD). Cells were analyzed under four conditions: 2 mM Ca^2+^, 10 mM EGTA, 2 mM Ca^2+^ with overexpressed CaM, and 10 mM EGTA with overexpressed CaM. Whole cell lysate (WCL) and biotinylated surface protein eluates (ELU) were probed with anti-Myc (Kv7.4), Na^+^/K^+^-ATPase (surface control), and GFP (cytosolic control) antibodies. **(B)** Quantification of surface Kv7.4 expression determined from the ratio of biotinylated Kv7.4 (Myc) to Na^+^/K^+^-ATPase in ELU samples. Values for each condition were normalized to the corresponding WT control. **(C)** Representative TIRF microscopy images of HEK293T cells expressing YFP-tagged Kv7.4-WT, TA and TD. **(D)** Quantification of the relative surface expression determined from TIRF image fluorescence intensities for WT, TA and TD. Data are presented as mean ± SD and statistical significance assessed by one-way ANOVA followed by Tukey’s multiple-comparison test.

Total internal reflection fluorescence (TIRF) microscopy experiments were conducted on HEK293T cells transfected with YFP-tagged Kv7.4-WT, TA and TD variants to further validate the surface expression findings. Micrographs showing YFP fluorescence of two representative cells from each variant are presented in **Figure 3C** with summary quantification in **Figure 3D**. Overall, the surface biotinylation and TIRF experiments show that the phospho-null and phosphomimetic variants do not significantly alter Kv7.4 channel surface expression.

### 2.3. Phosphorylation of Thr552 does not alter the secondary structure of free Kv7.4 B helix peptides in aqueous solution

To more precisely understand the mechanistic interplay between CaM, PIP_2_ and Kv7.4, we used independent peptides matching the A and B helix sequences in subsequent experiments. The peptide sequences are shown in **Figure 4A** below the Kv7.4 linear protein schematic. The **Q4S6JxA** peptide contains the cationic-rich S6Jx segment of the extended S6 transmembrane segment (highlighted in pink) followed by the A helix containing a classic IQ CaM-binding motif. The sequences for the B helix peptide include wild-type (**Q4Bx**), T552-phosphorylated (**Q4BpxT**), and a charge-neutralized KRK peptide containing K546A/R547A/K548A (**Q4BxAAA**), with Thr552 highlighted in green and the proposed PIP_2_ binding region highlighted in pink.

**Figure 4.**
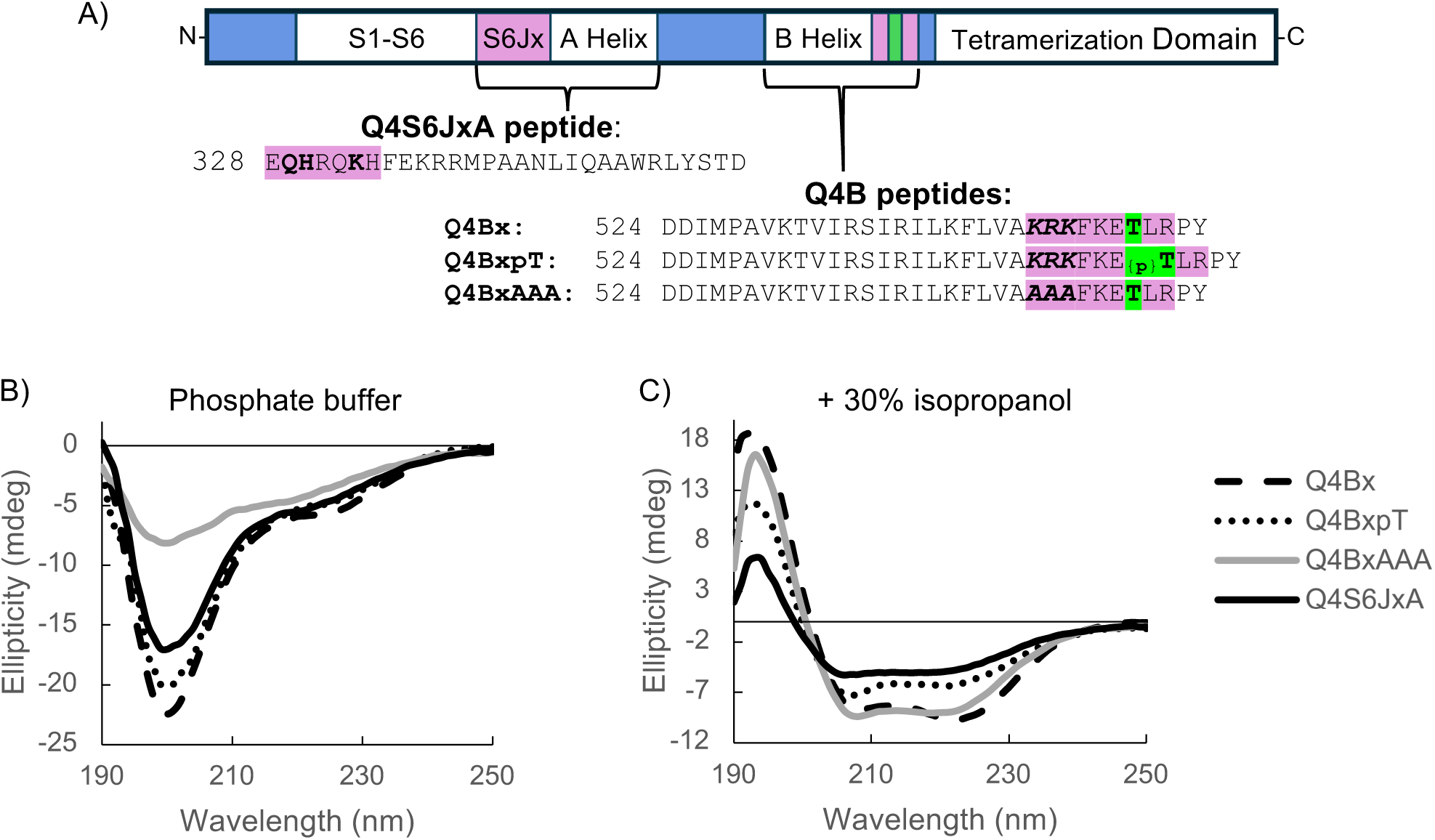
Circular dichroism spectra of Kv7.4 peptides. **(A)** Schematic showing the positions of the A and B helices within a linearized Kv7.4 subunit and the peptide sequences used in this study, based on UniProt entry P56696-1. Residues highlighted in pink indicate PIP_2_-binding regions outside the CaM-binding domains, and phosphorylation target Thr552 is highlighted in green. The polybasic KRK motif is shown in bold italics. **(B, C)** Circular dichroism spectra of the peptides Q4Bx (dashed line), Q4BxpT (dotted line), Q4BxAAA (gray line) and Q4S6JxA (black line) recorded in **(B)** aqueous buffer (5mM KH_2_PO_4_, 5mM NaCl, pH 7.4) or **(C)** the same buffer supplemented with 30% isopropanol.

Previously the A and B helices were reported to adopt a random coil or disordered structure in the absence of CaM^13^. Here, we examined whether inclusion of these adjacent regions, the S6Jx region in Q4S6JxA and the distal B region in Q4Bx, would affect the secondary structures of the independent peptides being used in this study. We performed circular dichroism spectroscopy to examine the secondary structures of these independent peptides. The data show that each of these peptides is largely unstructured (**Figure 4B** and **Table 1**), as determined using the ChiraKit online software^22^. This data suggest that inclusion of the adjacent regions or phosphorylation does not change the secondary structures of the helices.

**Table 1:** Predicted Secondary Structure Composition of Peptides.

| Peptide | Phosphate buffer |  |  | + 30% isopropanol |  |  |
| --- | --- | --- | --- | --- | --- | --- |
| | α- Helix | β-strand | Irregular | α- Helix | β- strand | Irregular |
| Q4Bx | 7.6% | 4.67% | 87.73% | 64.55% | 3.5% | 31.95% |
| Q4BxpT | 7.00% | 6.4% | 86.6% | 31.40% | 12.76% | 55.84% |
| Q4BxAAA | 9.61% | 15.87% | 74.53% | 35.49% | 8.64% | 55.87% |
| Q4S6JxA | 8.1% | 4.96% | 86.94% | 33.57% | 3.37% | 63.064% |

Isopropanol and other short-chain aliphatic alcohols are often used to dissolve hydrophobic synthetic peptides^23,24^. Alcohols are also reported to stabilize the α-helix content of peptides and proteins, as observed in β-lactoglobulin, poly (L-lysine), trypsin, and others^25–27^. When the peptides were dissolved in 30% isopropanol, they each exhibited prominent troughs at 208 and 222 nm, characteristic of α-helices (**Figure 4C** and **Table 1)**. While isopropanol increased the helical content of each peptide, both Q4BpxT and Q4BAAA still had approximately 55% disordered content and only ∼31-35% alpha helical content. In contrast, the unphosphorylated and wt Q4Bx peptide exhibited ∼32% disordered and ∼65% alpha helix content, consistent with the A helix and B helix having the propensity to adopt helical conformation. These data suggest that phosphorylation or a mutation in the KRK motif might lose some stability in their secondary structure.

### 2.4. Phosphorylation of Thr552 does not alter Ca^2+^/CaM interaction with the Kv7.4 B helix

Since phosphorylation at a similar site in KCNQ2 channels^12,28^ has been shown to attenuate the interaction between calmodulin and Kv7.2 channels, we questioned if the loss-of-function effect of the T552D mutation could be due to lack of binding of CaM to the channels. HSQC-NMR was performed on a 700 MHz Bruker NMR spectrometer to examine changes in ^15^N-^2^H-CaM spectra when bound to unphosphorylated (Q4Bx) vs the phosphorylated (Q4BpxT) peptide in the presence of Ca^2+^. **Figure 5A-C** shows the HSQC-NMR spectrum for Ca^2+^/CaM without peptide *(orange peaks)* compared to the spectra for Ca^2+^/CaM + Q4Bx *(turquoise peaks)* and Ca^2+^/CaM + Q4BxpT *(purple peaks).* We observed significant changes in the Ca^2+^/CaM spectrum after addition of each peptide, but the spectrum displayed by Ca^2+^/CaM + Q4Bx was nearly identical to that of Ca^2+^/CaM + Q4BxpT. The residues with peak height changes >2 standard deviations (SD) are shown in **Table ST2**. A total of 106 residues of Ca^2+^/CaM were significantly changed after the addition of Q4Bx, and 126 residues were changed by Q4BxpT, with 99 changed residues in common between the two spectra *(highlighted in cyan cells with dark blue font)*. These data show that both peptides caused significant changes to the N lobe, C lobe and linker of CaM in the presence of Ca^2+^. The residues that were changed on the CaM structure after adding peptide are colored in cyan or magenta on the PDB 1DMO structures (**Figures 5D & E)**. Based on this observation, phosphorylation of Thr552 would not cause major differences in the overall structure and interactions between Ca^2+^/CaM and the Kv7.4 B helix.

**Figure 5.**
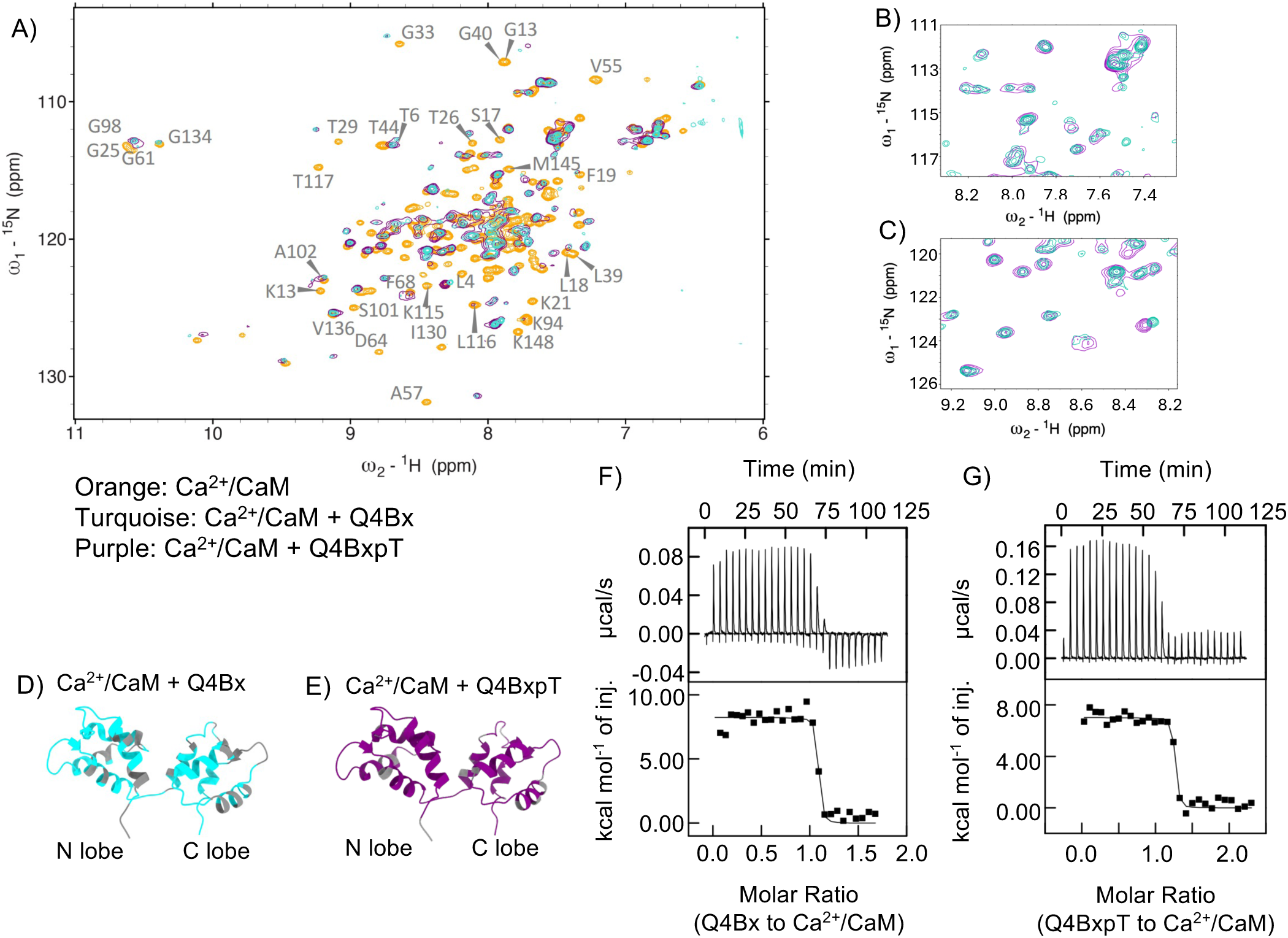
NMR and ITC shows that phospho-T552 does not alter the interactions and affinity between Ca^2+^/CaM and Kv7.4 B helix. **(A)** Overlay of of ^15^N-^2^H-Ca^2+^/CaM HSQC-NMR spectra acquired in the absence (orange peaks) or presence of Q4Bx *(turquoise)* or Q4BxpT *(purple).* Peptides were added at 1:1.2 CaM:peptide molar ratio**. (B, C)** Expanded regions of the overlaid HSQC spectra showing extensive peak overlap between Ca^2+^/CaM bound to Q4Bx and Q4BpxT. **(D, E)** Structure of Ca^2+^/ CaM structure (PDB: 1DMO) highlighting residues that exhibited chemical shift perturbations and/or change in peak height (>2 SD) upon peptide binding. **(F, G)** Representative ITC isotherms of Q4Bx (F) or Q4BpxT **(G)** into Ca^2+^/CaM. Peptide concentrations ranged from 35 - 60 μM and were titrated into 5 μM Ca^2+^/CaM at 20^∘^C.

To validate this finding, we used isothermal titration calorimetry (ITC) to determine the affinity of Ca^2+^/CaM with Q4Bx **(Figure 5F)** and Q4BxpT **(Figure 5G)** peptides. We determined the K_d_ ∼0.95 nM for Q4Bx and K_d_ ∼1.6 nM for Q4BxpT binding to Ca^2+^/CaM, with no significant difference between the two values **(Table 2)**. This result is slightly higher affinity than our previously reported K_d_ value for the shorter Q4B peptide of ∼4 nM^13^, suggesting that the additional residues added to the distal B helix peptides strengthen the interactions with Ca^2+^/CaM. These data suggest that phosphorylation of Thr552 does not significantly change the affinity or alter the interactions between Ca^2+^/CaM and the Kv7.4 B helix.

**Table 2.** Summary of ITC results of Ca^2+^/CaM binding the Q4Bx and Q4BxpT peptides.

| Peptide | n<br>(# trials) | K<sub>a</sub><br>(M<sup>-1</sup>) | K<sub>d</sub><br>(nM) | N<br>(stoichiometry) | $\Delta H$<br>(kcal/mol) | $\Delta S$<br>(cal/mol/deg) |
| --- | --- | --- | --- | --- | --- | --- |
| Q4Bx | 3 | 1.10e09 ± 2.76e08 | 0.95 ± 0.23 | 1.15 ± 0.10 | 7.88 ± 0.05 | 68.2 ± 1.37 |
| Q4BxpT | 3 | 6.42e08 ± 1.85e08 | 1.67 ± 0.57 | 1.08 ± 0.13 | 6.42 ± 0.06 | 62.1 ± 6.56 |

Values are mean ± S.D.

### 2.5. Phosphorylation of Thr552 of the Kv7.4 B helix induces domain-specific line broadening in the apoCaM N-lobe

HSQC-NMR was also used to probe the effects of phosphorylated Thr552 on the interactions between apoCaM and the B helix. The spectrum of ^15^N-^2^H-apoCaM was compared in the presence of Q4Bx or Q4BxpT peptides. **Figure 6A** shows the spectrum of apoCaM without peptide (*orange peaks*) overlaid with apoCaM + Q4Bx *(pink peaks)* and apoCaM + Q4BxpT *(green peaks)*. We observed more green peaks (apoCaM + Q4BxpT) overlap the orange peaks (apoCaM) than do the pink peaks (apoCaM + Q4Bx). The expanded views in **Figure 6B-C** show distinct changes in apoCaM induced by Q4Bx compared to Q4BxpT. The apoCaM residues that exhibited a significant shift or line broadening were determined by calculating changes in the spectra >2 SD of the mean peak height **(Table ST2)**, revealing that Q4Bx caused changes to 65 different residues throughout the CaM protein, and colored *pink* in **Figure 6D**. In contrast, the addition of Q4BxpT caused significant changes to only 20 residues of apoCaM, all of which localized only to the C lobe, indicated in *green* in **Figure 6E**. **Figure 6F**, *(orange),* shows that 19 of the same apoCaM C lobe residues interact with both phosphorylated and unphosphorylated B helix. The localized signal attenuation indicates that the C lobe selectively enters an intermediate exchange regime on the NMR timescale, ruling out non-specific global aggregation or precipitation, and demonstrating a domain-specific responsiveness to the B helix peptide. The NMR data suggest that phospho-Thr552 reorganizes the apoCaM-B helix interaction.

**Figure 6.**
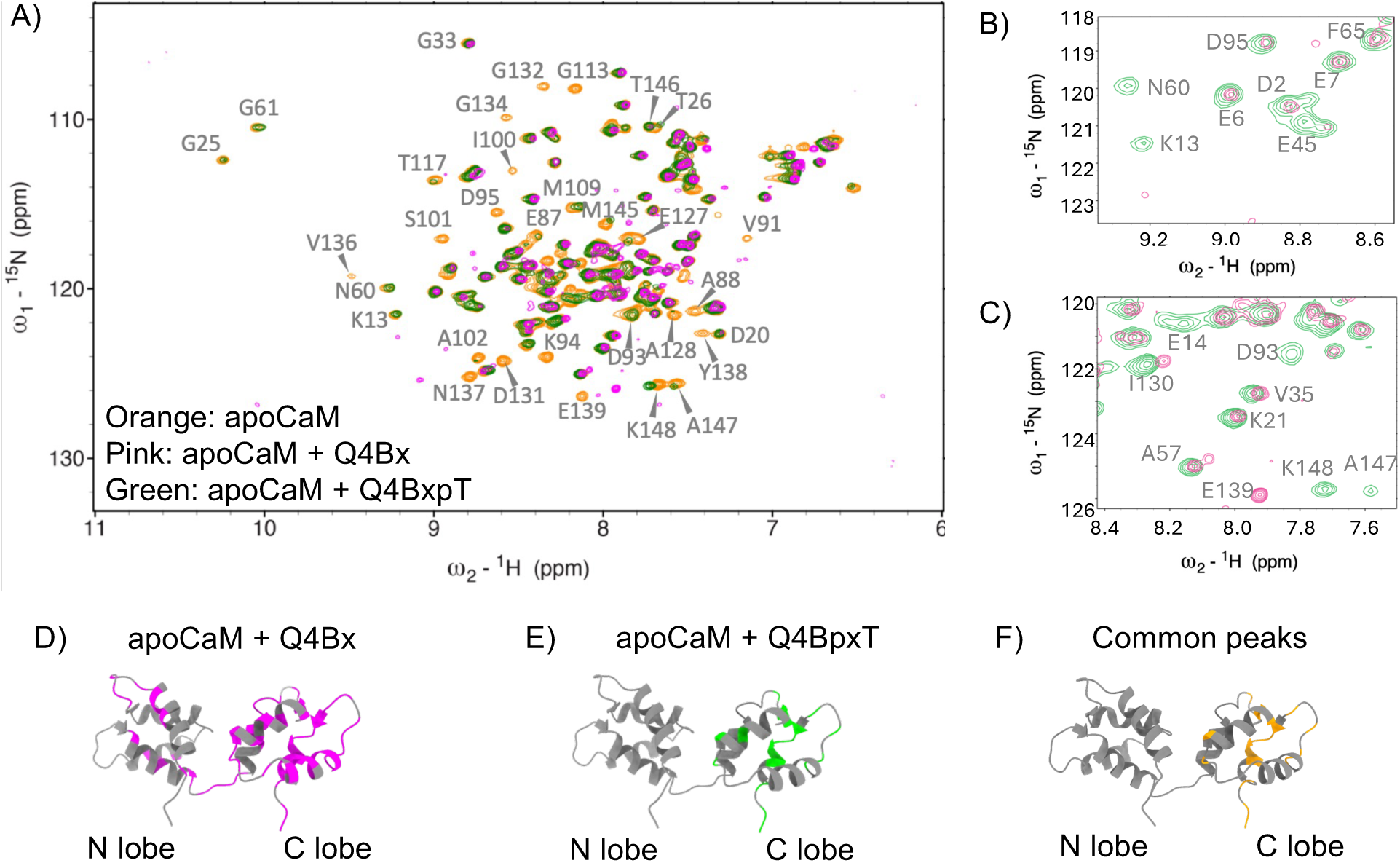
HSQC-NMR analysis reveals reduced interactions between apoCaM and the phosphorylated Kv7.4 B helix. **(A)** Overlay of of ^15^N-^2^H-apoCaM HSQC-NMR spectra acquired in the absence (orange peaks) or presence of Q4Bx *(pink)* or Q4BxpT *(green).* Peptides were added at 1:1.2 CaM:peptide molar ratio. **(B, C)** Expanded regions of the overlaid HSQC spectra highlighting spectral differences between apoCaM bound to Q4Bx and Q4BpxT. **(D-F)** Solution NMR structure of apoCaM (PDB: 1DMO) highlighting residues that exhibited chemical shift perturbations and/or change in peak height (>2 SD) upon peptide binding. Residues affected by Q4Bx are shown in pink **(D),** Residues affected by BpxT are show in green **(E),** and residues affected by both peptides are shown in orange **(F).**

### 2.6. Phosphorylation of Thr552 remodels the local interaction surrounding Lys548 without disrupting the CaM-Q4A-B complex

To investigate how phosphorylated Thr552 influences structural organization of the Kv7.4 regulatory domain complex with CaM, we performed molecular dynamics (MD) simulations using the crystal structure of the apoCaM-Q4AB (PDB: 6B8L) (**Figure 7A**). Unphosphorylated and phosphorylated complexes were simulated in both apoCaM and Ca^2+^/CaM conditions. All systems reached a stable RMSD plateau during the simulation period of 50 ns, indicating structural equilibrium and allowing comparison of phosphorylation-dependent conformational changes (**Figure S3**).

**Figure 7.**
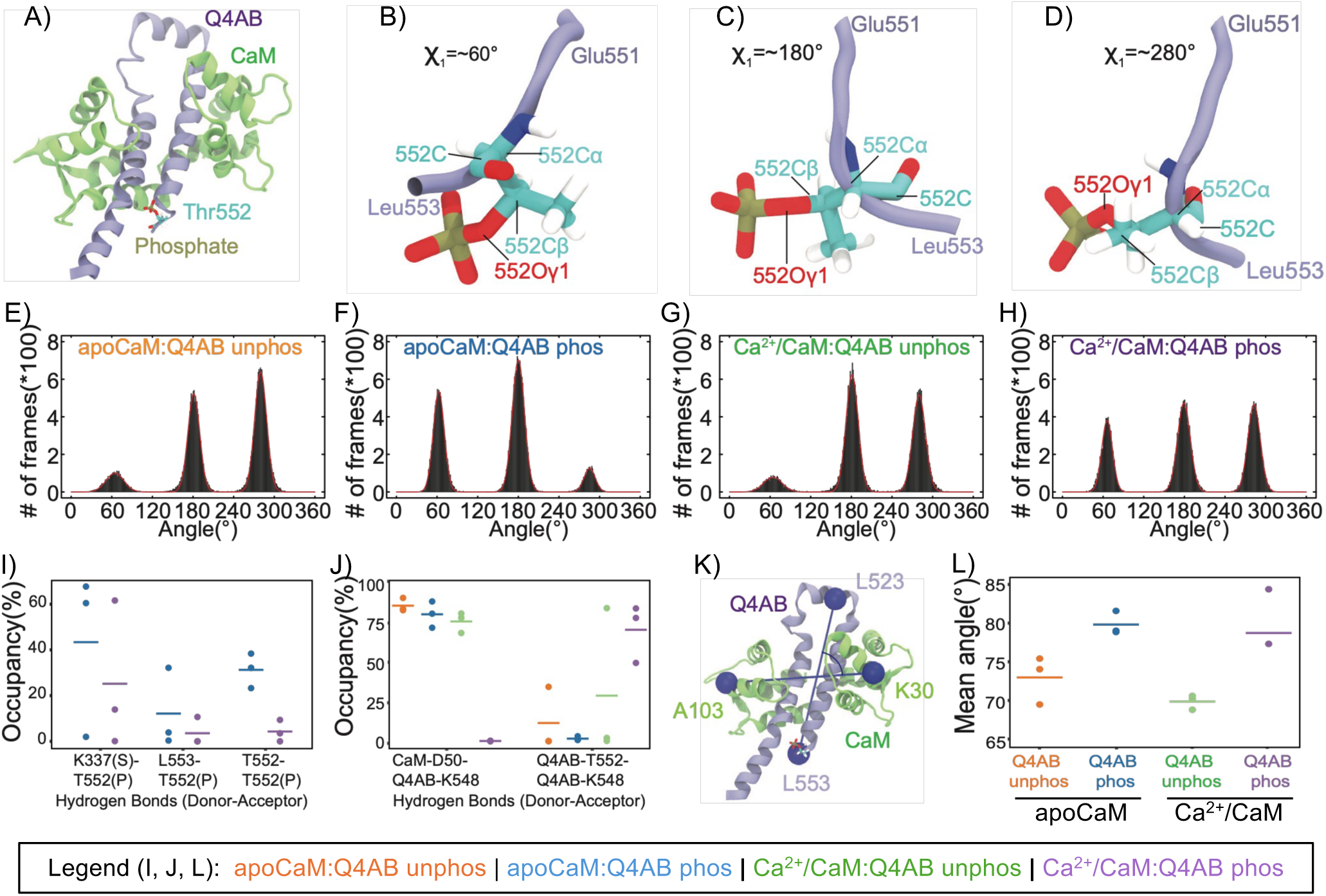
Molecular dynamics simulations reveal phosphorylation-induced local structural perturbations in the CaM:Q4AB complex under ApoCaM and Ca^2+^/CaM conditions. **(A)** Representative structure of the phosphorylated CaM:Q4AB complex (PDB: 6B8L). CaM is shown in green, Q4AB in blue, and the phosphate group is colored by atom type. The unphosphorylated system lacks the modeled phosphate group. **(B-D)** Enlarged views of the phosphorylated Thr552 region corresponding to C–Cα–Cβ–Oγ1 dihedral angles of 60°, 180°, and 280°, respectively. **(E-H)** Distribution of C–Cα–Cβ–Oγ1 dihedral angle populations for the apoCaM:Q4AB unphosphorylated **(E),** apoCaM:Q4AB phosphorylated (F), Ca^2+^/CaM:Q4AB unphosphorylated **(G),** and Ca^2+^/CaM:Q4AB phosphorylated (H) systems. (I) Occupancy of hydrogen bonds formed by phosphate oxygens in Thr552-phosphorylated apoCaM:Q4AB and Ca^2+^/CaM:Q4AB complexes. (S) denotes an interacting sidechain amide hydrogen, and (P) denotes the interacting oxygen of phosphate group. No notation indicates a backbone interaction. **(J)** Change in hydrogen bond occupancies across all systems. **(K)** Definition of the reference vectors used to quantify the CaM:Q4AB interaction. One vector connects the Cα atoms of CaM residues Ala104 and Lys31, whereas the second vector connects the Cα atoms of Leu523 and Leu558 of Q4AB. **(L)** Conformational displacements between CaM and Q4AB in the four simulation systems.

We first examined the side-chain orientation of Thr552 by monitoring its χ1 dihedral angle throughout the trajectory (**Figure 7 B-H**). Thr552 occupied three major rotameric states centered around approximately 60°, 180°, and 280°. In the unphosphorylated systems for both apoCaM and Ca^2+^/CaM conditions, the Thr552 side chain predominantly populated orientations of ∼180° and ∼280°. Phosphorylation altered distribution among these conformers in both apoCaM and Ca^2+^/CaM systems, with the distribution shifted to a substantially increased χ_1_ value of ∼60° and reduced occupancy of the ∼280° state. This shift was most pronounced in the phosphorylated apoCaM-Q4AB complex and suggests that addition of the phosphate group produces a preferred orientation of the distal B side chain that is determined by the presence of Ca^2+^.

To understand the structural consequence of this reorientation, we examined the hydrogen bond networks surrounding Thr552. Phosphorylation promoted new intramolecular hydrogen bonds with the phosphate group and neighboring residues, including the side-chain amide hydrogen of Lys337 and the backbone amide hydrogen of Leu553 on the B helix (**Figure 7I**). These interactions stabilized local conformations around the phosphorylation site and redistributed nearby intermolecular contacts. Notably, Lys548, located within the polybasic region of the B helix, exhibited phosphorylation-dependent changes in its interaction network (**Figure 7J**). In the unphosphorylated complex, Lys548 participated in intermolecular interactions with Asp50 of the CaM N-lobe, consistent with the experimental NMR results showing the Asp50 peak as one of the significantly shifted peaks with of apoCaM + unphosphorylated B helix while that same residue for apoCaM + phosphorylated B helix was unaffected **(Figure 6, Table ST2).** Following phosphorylation, occupancy of these interactions decreased substantially and was abolished in the Ca^2+^/CaM phosphorylated simulations (*purple circles*) in **Figure 7J**. Concurrently, the phosphate group of Thr552 formed new local interactions that altered the orientation of neighboring residues, forming new hydrogen bonds with Lys 548. These findings suggest that phosphorylation does not disrupt the overall CaM-Q4AB complex but instead reorganizes local contacts involving distal B helix KRK motif.

We also examined the effects of phosphorylation on the overall changes in secondary structure and global structure of the CaM-Q4AB complex. Ramachandran plot analysis **(Figure S4)** indicated significant changes in secondary structure of only the N terminal residue positions 4-7 of CaM which are likely flexible in nature and do not form direct contacts with the B helix. Conformational displacement calculations of the CaM-Q4AB interface show an increased angle between vectors representing the horizontal axis of CaM and the vertical axis of the phosphorylated Q4AB B-helix (**Figure 7L**), indicating a repositioning of the complex upon phosphorylation. These data along with the apoCaM NMR experiments support phosphorylation impacting the global structure of the CaM–Q4AB complexes.

Overall, the MD simulations indicate that phosphorylation of Thr552 induces local structural rearrangements within the distal B helix rather than global disruption of the CaM-Q4AB complex. The most prominent change involves remodeling of interactions of surrounding Lys548, a residue positioned within the conserved polybasic KRK motif. These results provide a structural framework linking Thr552 phosphorylation to altered regulation of the distal B helix and motivated subsequent experiments examining the distal KRK motif.

### 2.7. Phosphorylation of Thr552 does not substantially disrupt apoCaM association with the Kv7.4 B helix

To determine the affinity of apoCaM to Q4Bx (wt) and Q4BxpT (phospho-T552) peptides, we performed microscale thermophoresis (MST) experiments using CaM conjugated with Red-NHS dye in ChHBS buffer **(Figure 8A-D)**, as we were unable to obtain binding data for apoCaM-B helix using ITC, likely because of their lower binding affinities. Control experiments are shown in **Figure S5**. The dissociation constant for Q4Bx-apoCaM was approximately 10 µM (95% CI, 7-15 µM, **Figure 8A)** similar to values previously reported for apoCaM binding a shorter version of the Q4B peptide lacking Thr552^13^. Phosphorylation of Thr552 produced a modest increase in apoCaM affinity, with Q4BxpT exhibiting a K_d_ ∼4.6 µM (95% CI of 2.8-7.4 µM), **(Figure 8B, E)**. These results are summarized in **Table 3**. Despite this statistically significant difference, both peptides bound apoCaM with affinities in the same low-micromolar range, indicating that phosphorylation does not substantially alter overall apoCaM association with the Kv7.4 B helix. Consequently, phosphorylation-dependent changes in apoCaM interactions are unlikely to arise from loss of binding affinity and instead may reflect changes in the mode of interaction observed by NMR **(Figure 6)**

**Figure 8.**
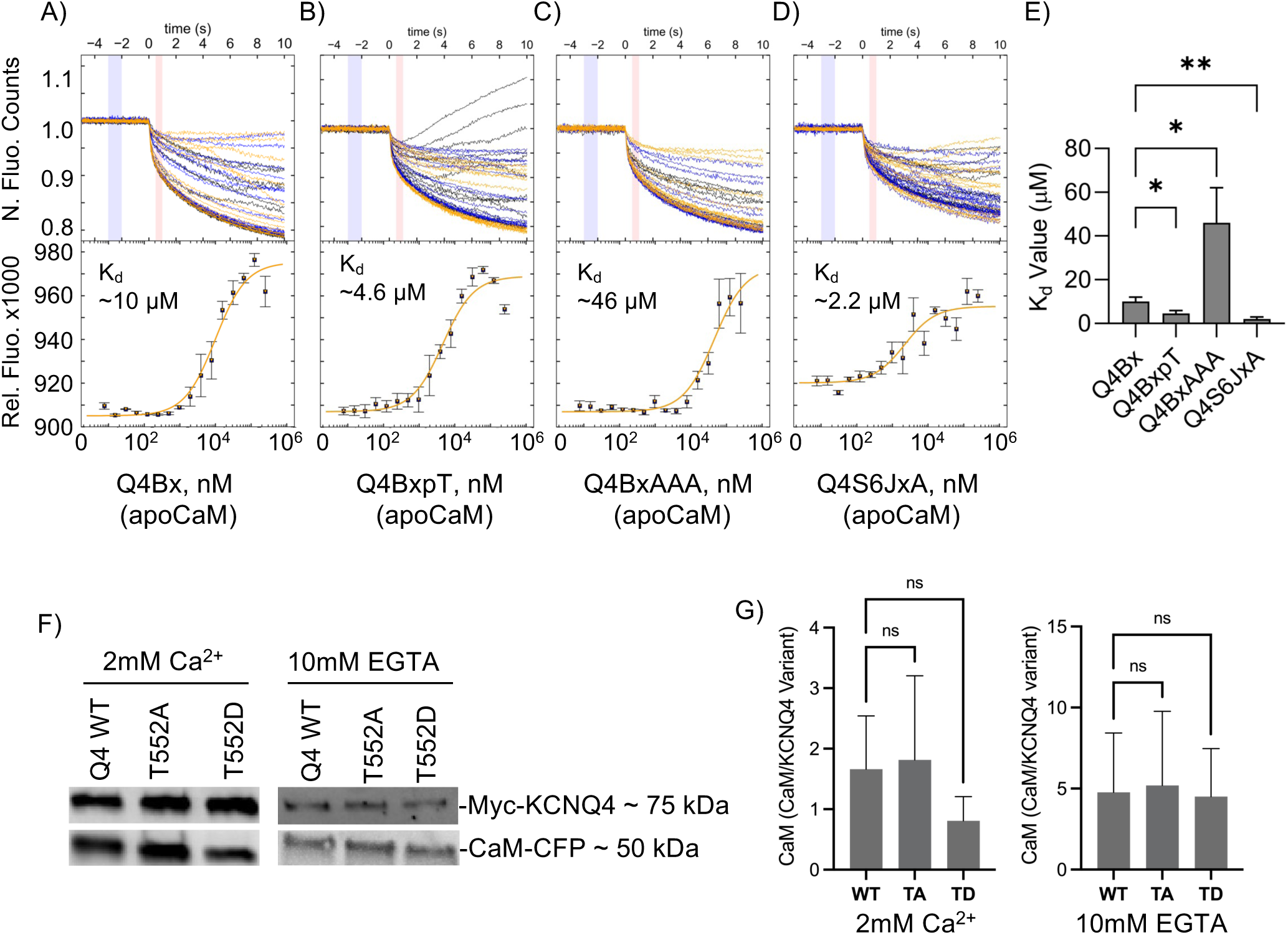
Binding affinities between apoCaM-Q4 peptides. (A-D) Representative MST thermophoresis traces and binding curves for Q4Bx **(A)**, Q4BxpT **(B),** Q4BxAAA **(C),** and Q4S6JxA **(D)** titrated against 50 nM of Red-NHS-labeled apoCaM in HBS supplemented with 0.1 mM EGTA, 0.05% Tween-20 (pH 7.4). **(E)** Summary of equilibrium dissociation constants K_d_ obtained from MST measurements and analyzed using PALMIST. **(F)** Representative figures of co-immunoprecipitation of Myc-tagged Kv7.4 WT, T552A, and T552D with CaM-CFP under elevated Ca^2+^ (left) and EGTA (right) conditions. Similar amounts of CaM-CFP were co-immunoprecipitated with all three Kv7.4 variants across three independent experiments. **(G)** Quantification of the co-IP results for Ca^2+^ (left) and EGTA (right) conditions. Data for all experiments are presented as mean ± SD from n=3 independent experiments. Statistical significance was assessed using unpaired *t* tests.

**Table 3:** Binding affinities of Q4 peptides with apoCaM.

| Peptide | K<sub>d</sub> ($\mu\text{M}$) | 95% CI ($\mu\text{M}$) | SD ($\mu\text{M}$) |
| --- | --- | --- | --- |
| Q4Bx | 10 | [7.0, 15] | $\pm 2.0$ |
| Q4BxpT | 4.6 | [2.8, 7.4] | $\pm 1.3$ |
| Q4BxAAA | 46 | [28, 80] | $\pm 16$ |
| Q4S6JxA | 2.2 | [0.9, 6.6] | $\pm 1.1$ |

Data represent n=3 experiments. K<sub>d</sub> values were calculated and reported through PALMIST and analyzed in GraphPad Prism 10.

To validate this result, we performed co-immunoprecipitation of the myc-tagged Kv7.4 wt, TA and TD with overexpressed CaM-CFP in HEK293T cells in the presence and absence of Ca^2+^. CaM was co-immunoprecipitated with all variants in the presence and absence of Ca^2+^ **(Figure 8F**). Quantification of the co-immunoprecipitated CaM revealed no significant differences among WT, T552A and T552D channels **(Figure 8G)**. These results further support the conclusion that phosphorylation of Thr552 does not promote complete dissociation of the CaM-Kv7.4 complex. However, co-IP cannot resolve the domain-specific changes detected by NMR and MD simulations which indicate that phosphorylation alters the mode of apoCaM interaction without substantially affecting overall association.

MST also revealed a significant reduction of affinity (approximately 4.5X) between the Q4BxAAA peptide and apoCaM (K_d_ ∼46 µM) **(Figure 8C)**, reinforcing the MD results that identified Lys548 as an important contributor to apoCaM interactions with the B helix. The Q4SJxA peptide bound apoCaM with a K_d_ of ∼2.2 µM as determined by MST (**Figure 8D**). Interestingly, the Q4S6JxA-apoCaM interaction is much stronger than the binding affinity of the short Q4A peptide binding to apoCaM (K_d_ ∼45 µM) **(Figure S6)** which was beyond the detection limit in our earlier study^13^. This result demonstrates that the S6Jx region is critical for the A helix to fully engage with apoCaM.

Together, these results support an important role for the distal B helix KRK motif in apoCaM interactions and indicate that phosphorylation of Thr552 alters, but does not abolish, apoCaM association with the Kv7.4 regulatory domain.

### 2.8. Phosphorylation of Thr552 reduces PIP_2_ interactions with the Kv7.4 regulatory domain

Because Lys548 resides within a polybasic region (KRK motif) previously implicated in CaM interactions^13^ and proposed to contribute to PIP_2_ recognition, we reasoned that phosphorylation-dependent remodeling of this region could influence lipid binding without requiring dissociation of CaM. Consistent with this possibility, our MD simulations identified phosphorylation-dependent changes in the local interaction network surrounding Lys548 **(Figure 7J)**. To determine whether phosphorylation alters accessibility or function of this cationic binding surface, we next examined PIP_2_ interactions with the Kv7.4 regulatory domain.

PIP_2_ interactions were examined using the fluorescent PIP_2_ headgroup analog PHG-FL in a fluorescence polarization assay **(Figure 9A, B, Table 4)**. In this analog, the fluorescein tag is conjugated to carbon 1 in place of the bond that connects to the fatty acyl chains, preserving the phosphoryl groups at carbons 4 and 5 to interact with PIP_2_ binding motifs. As expected, PHG-FL exhibited no measurable interaction with CaM alone in either the presence or absence of Ca^2+^ (**Figure 9A**, *grey and dotted black lines & symbols*). The molecular weights of the Q4A and B helix peptides alone are too small to increase the polarized fluorescence during binding, but if PHG-FL binds to them while they are pre-bound to CaM then the fluorescence is expected to increase. We observed high fluorescence polarization values for PHG-FL binding the Q4Bx + CaM complex in the presence and absence of Ca^2+^ (**Figure 9A**). Titrating the complexes against the PIP_2_ analog allowed us to calculate the affinities at K_d_ ∼14 µM for Q4Bx + Ca^2+^/CaM and K_d_ ∼22 µM for Q4Bx + apoCaM, with no significant difference between these two results (**Figure 9B, and Table 4)**. Because the isolated peptides are below the threshold required for detection by fluorescence polarization, PIP_2_ binding was assessed using peptide-CaM complexes. The resulting increase in fluorescence polarization is consistent with simultaneous association of CaM and PIP_2_ with the Kv7.4 regulatory domain. These results show strong agreement with the MST results in **Figure 9C**, with PHG-Cy5, a similar analog to PHG-FL just with a different tag. PHG-Cy5 interacted with the B helix with a K_d_ ∼4.5 µM. Together, these findings indicate that the distal B helix contributes to PIP_2_ binding in both apoCaM- and Ca^2+^/CaM-associated complexes.

**Figure 9.**
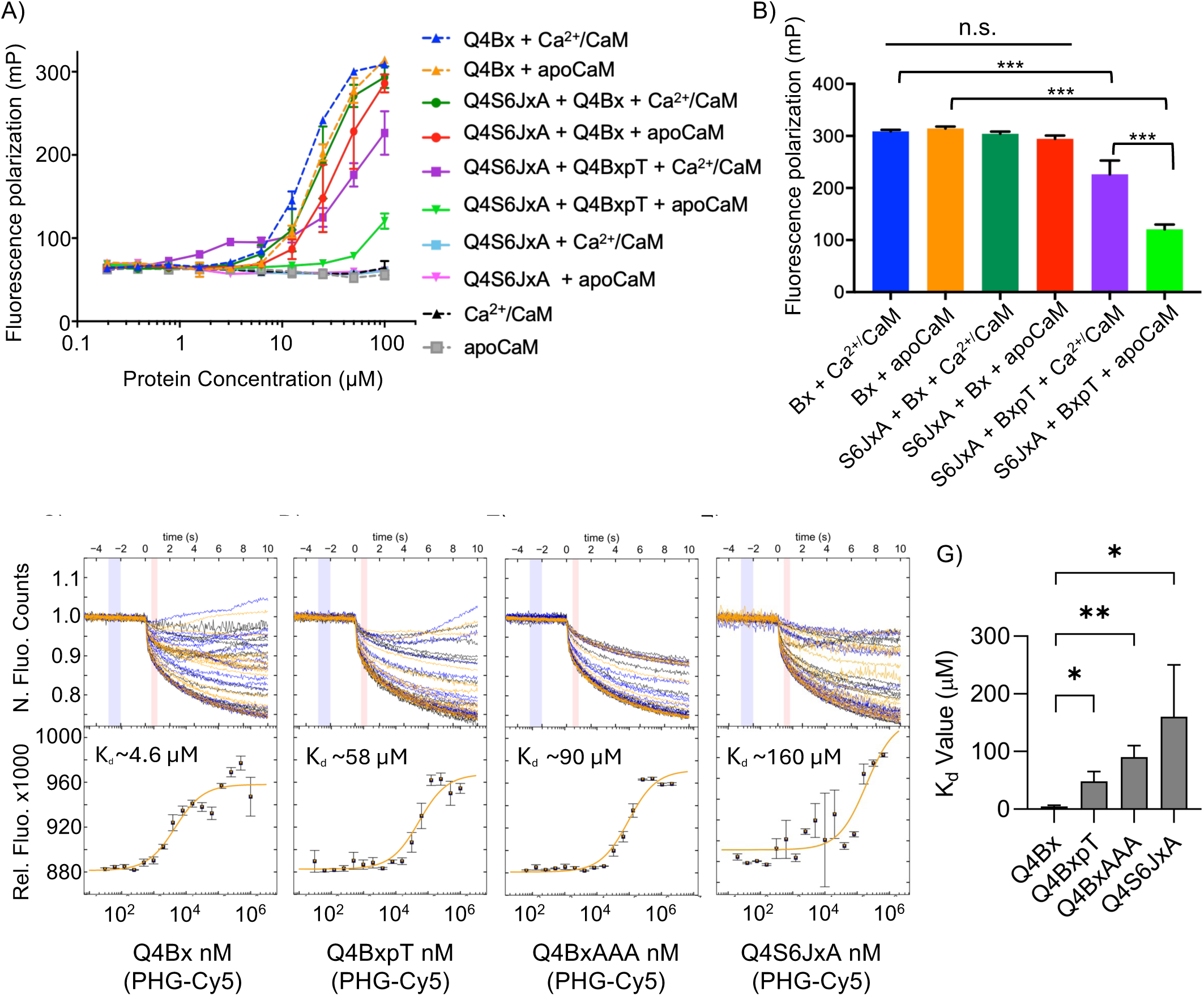
Binding affinities of the PIP_2_ headgroup for the Kv7.4 A and B helix peptides. **(A)** Fluorescence polarization binding curves obtained by titrating twofold serial dilutions of CaM-peptide complexes against 50 nM fluorescein-labeled PIP_2_ headgroup (PHG-FL) under Ca^2+^-free and Ca^2+^-bound conditions. **(B)** Summary of fluorescence polarization values measured at 100 μM protein complex concentration. Colors correspond to the peptide combinations shown in **(A)**. (**C-F)** MST thermophoresis traces and binding curves for Q4Bx **(C)**, BxpT **(D)**, BxAAA **(E)**, S6JxA **(F)** titrated against 50 nM of PHG-Cy5 in HBS supplemented with 0.1 mM EGTA, 0.05% Tween-20 (pH 7.4). **(G)** Summary of equilibrium dissociation constants K_d_ obtained from MST measurements and analyzed using PALMIST. Data are presented as mean ± SD. MST experiments represent n=3 independent experiments whereas fluorescence polarization measurements represent n=2 independent experiments. Fluorescence polarization data were analyzed by one-way ANOVA followed by Tukey’s multiple-comparison test and MST-derived K_d_ values were compared using unpaired t tests.

**Table 4:**
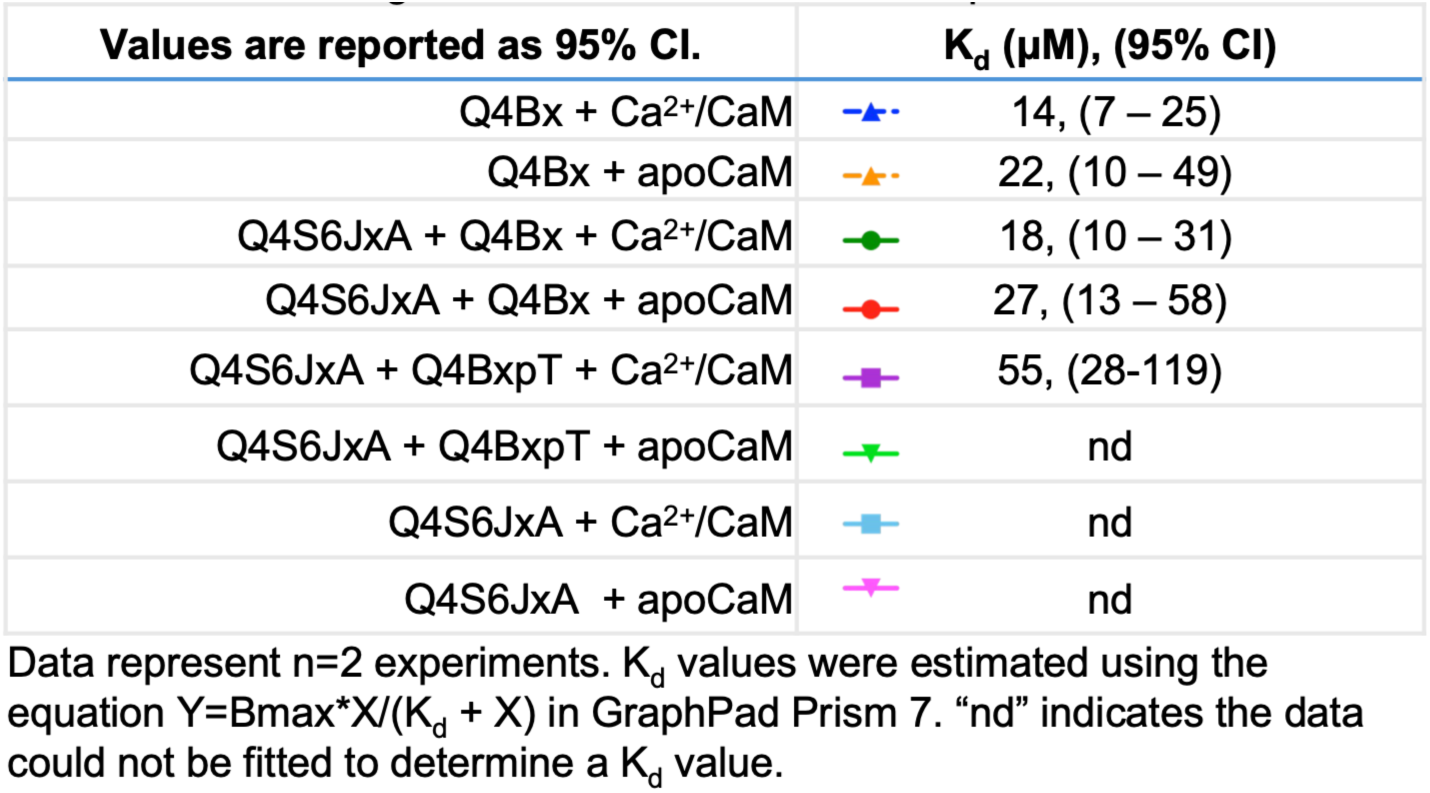
PIP2 binding affinities for KCNQ4 A+B complex with CaM.

| Values are reported as 95% CI. | $K_d$ (μM), (95% CI) | |
| --- | --- | --- |
| Q4Bx + Ca<sup>2+</sup>/CaM |  | 14, (7 – 25) |
| Q4Bx + apoCaM |  | 22, (10 – 49) |
| Q4S6JxA + Q4Bx + Ca<sup>2+</sup>/CaM |  | 18, (10 – 31) |
| Q4S6JxA + Q4Bx + apoCaM |  | 27, (13 – 58) |
| Q4S6JxA + Q4BxpT + Ca<sup>2+</sup>/CaM |  | 55, (28-119) |
| Q4S6JxA + Q4BxpT + apoCaM |  | nd |
| Q4S6JxA + Ca<sup>2+</sup>/CaM |  | nd |
| Q4S6JxA + apoCaM |  | nd |

Data represent n=2 experiments. $K_d$ values were estimated using the equation $Y=B_{max} \cdot X / (K_d + X)$ in GraphPad Prism 7. “nd” indicates the data could not be fitted to determine a $K_d$ value.

**Table 5:** Binding affinites of Q4 peptides with PHG-Cy5 without Ca^2+^.

| Peptide | $K_d$ (μM) | 95% CI (μM) | SD (μM) |
| --- | --- | --- | --- |
| Q4Bx | 4.6 | [2.0, 11] | ±1.9 |
| Q4BxpT | 48 | [28, 83] | ±17 |
| Q4BxAAA | 90 | [60, 140] | ±20 |
| Q4S6JxA | 160 | [70, 470] | ±90 |

Data represent n=3 experiments. $K_d$ values were calculated and reported through PALMIST and analyzed in GraphPad Prism 10.

Unexpectedly, the CaM-Q4S6JxA complexes exhibited little detectable PIP_2_ binding despite containing residues as identified as PIP_2_-interacting in cryo-EM structures (**Figure 9A**, *light blue and pink lines & symbols)*. This observation corroborates with our MST findings, in which the binding affinity of Q4S6JxA to PHG-Cy5 is also low K_d_ ∼160 µM (**Figure 9F**). Since PIP_2_ contacts this region in cryo-EM structures **(Figure 1A**), the weak binding observed here may indicate that the isolated Q4S6JxA peptide lacks the structural context required for efficient PIP_2_ recognition or that CaM occludes residues contributing to this interaction.

The triple complex of S6JxA+Q4Bx+CaM exhibited similar affinity to PIP_2_ as observed with the complex of Q4Bx + CaM. Under high [Ca^2+^] the affinity of this triple complex for the PHG-FL ligand was K_d_ ∼18 µM, (**Figure 9A**, *dark green line and circles)*, and in the absence of Ca^2+^ the affinity was K_d_ ∼27 µM, *(red line and circles)*. These results were not significantly different from each other **(Figure 9A**, *dark green and red bars*) and are consistent with PIP_2_ interacting with the cationic residues surrounding the A and B helices while bound to CaM.

We used the Q4BxpT peptide to examine how phosphorylated Thr552 impacts the interactions between PIP_2_ and the CaM+A+B complexes. S6JxA + Q4BxpT + Ca^2+^/CaM complex *(purple lines & symbols)* exhibited significantly reduced PHG-FL fluorescence polarization compared to Q4Bx + Ca^2+^/CaM *(dotted blue line & triangles)* (**Figure 9A-B**, **Table 4**). In contrast, while the fluorescence of PHG-FL began to increase with higher concentration of the S6JxA + Q4BxpT + apoCaM complex *(lime green line & symbols)*, its fluorescence polarization levels were too low to determine a K_d_ value (**Figure 9A**). There was a significant difference between these two results at 100 µM protein **(Figure 9B)**. Additionally, the PIP_2_ fluorescence polarization of the S6JxA + Q4BxpT + CaM complexes were significantly decreased compared to the S6JxA + Q4Bx + CaM complexes in the presence and absence of Ca^2+^ (**Figure 9B**, **Table 4**). This result is consistent with the MST measurements of Q4BpxT which exhibited substantially lower affinity (K_d_ ∼58 µM) for PHG-Cy5 than the unphosphorylated peptide (**Figure 9D).** These findings demonstrate that phosphorylated Thr552 substantially reduces PIP_2_ interactions with the CaM-associated Kv7.4 regulatory domain.

**Figure 9E** shows the binding affinity between the PIP_2_ headgroup and Q4BxAAA peptide is strongly reduced, with K_d_ ∼90 µM. Since there is a significant reduction in binding affinities between BxAAA to PHG-Cy5 compared to Bx to PHG-Cy5, this suggests that PHG-Cy5 can bind to the B helix at the KRK region, which could compete with apoCaM for binding as the KRK is a known binding site for apoCaM^13^. This result strongly supports the KRK region on the distal B helix as a PIP_2_ binding site.

Together, the fluorescence polarization and MST experiments demonstrate that the distal B helix KRK motif contributes directly to PIP_2_ recognition and that phosphorylation of Thr552 markedly weakens these interactions. These findings provide a mechanistic explanation for the loss-of-function phenotype observed in the T552D mutant and support a model in which phosphorylation primarily regulates Kv7.4 activity by reducing PIP_2_ binding rather than loss of CaM association.

## DISCUSSION

In this study, we identified Thr552 as a phosphorylation-sensitive regulatory site that negatively regulates Kv7.4 activity. Phosphomimetic substitution of Thr552 produced a pronounced loss-of-function phenotype characterized by reduced current density and a depolarizing shift in voltage-dependent activation. These effects were observed under both high- and low-calcium conditions and persisted despite CaM overexpression, indicating a mechanism distinct from classical calcium-dependent suppression of Kv7 channels. The loss-of-function phenotype was not accompanied by altered surface expression, indicating that phosphorylation predominantly affects channel regulation and gating rather than trafficking.

Although Thr552 is positioned near the distal end of the B helix, within the CaM-Kv7 regulatory interface, multiple approaches in this study demonstrated that phosphorylation does not substantially disrupt CaM association. ITC revealed minimal effects on Ca^2+^/CaM binding, MST showed only modest changes in apoCaM affinity, and co-immunoprecipitation demonstrated comparable association of CaM with WT, T552A, and T552D channels. Together, these findings argue against a model in which phosphorylation inhibits Kv7.4 through dissociation of the CaM-channel complex.

While overall CaM association was preserved, NMR spectroscopy revealed that phosphorylation of Thr552 produced a marked redistribution of NMR perturbations, with residues in the C lobe remaining responsive whereas N-lobe perturbations were substantially reduced. Consistent with these observations, molecular dynamics simulations identified remodeling of the local interaction network surrounding Lys548 within the distal B-helix polybasic KRK motif. One interpretation is that the unphosphorylated B helix engages both lobes of apoCaM, whereas phosphorylation shifts the interaction toward a C-lobe-dominated binding mode. Although the present data do not define the precise orientation of apoCaM on the B helix, they suggest that phosphorylation alters lobe-specific contacts without abolishing overall association. These observations raise the possibility that the apoCaM N-lobe contributes to interactions with more distal regions of the B helix under low-Ca^2+^ conditions and that these contacts are destabilized following phosphorylation of Thr552. Notably, existing cryo-EM and crystal structures place the CaM N lobe adjacent to the B helix and the C lobe adjacent to the A helix, supporting the idea that phosphorylation may selectively perturb a subset of CaM-B helix interactions while preserving overall complex formation^9,13,20^. Together, these findings support a model in which phosphorylation reorganizes CaM interactions without abolishing binding.

In contrast to the relatively modest effects on CaM association, phosphorylation produced a marked reduction in PIP_2_ binding. Both fluorescence polarization and MST measurements demonstrated reduced interactions between phosphorylated Kv7.4-derived peptides and PIP_2_ analogs. Mutation of the KRK motif similarly impaired PIP_2_ binding and, together with the altered Lys548 interactions observed in the MD simulations, identified the distal B-helix polybasic region as an important determinant of lipid recognition. These observations indicate that phosphorylation of Thr552 primarily acts by weakening PIP_2_ interactions with the Kv7.4 regulatory domain.

Our findings are broadly consistent with previous studies implicating phosphorylation of the conserved distal B-helix serine/threonine residue in regulation of Kv7 channel activity through PIP_2_-dependent mechanisms. In Kv7.2, phosphorylation of the conserved B helix serine residue homologous to Kv7.4 Thr552 alters channel sensitivity to receptor-mediated inhibition and PIP_2_ depletion^29^. Hoshi et al. reported that mutation of this residue reduced muscarinic suppression of channel activity, whereas subsequent studies from that lab demonstrated that phosphomimetic substitution increased channel sensitivity to PIP_2_ depletion and proposed a mechanism involving altered CaM-dependent regulation^12,28^. Similarly, studies of Kv7.4 reported altered PIP_2_-dependent regulation following mutation Thr552 (Thr553 in that study) modified channel responses associated with PIP_2_ signaling^30^. Our results extend these observations by providing direct biochemical evidence that phosphorylation weakens PIP_2_ interactions with the Kv7.4 regulatory domain. The distal B-helix KRK motif appears to represent a critical regulatory node because it contributes to both CaM and PIP_2_ interactions. Consistent with this interpretation, Tobelaim et al. identified the analogous region in Kv7.1 as a site where CaM and PIP_2_ exert competing regulatory influences. Our MST and fluorescence polarization experiments support a similar role for the Kv7.4 KRK motif, while additionally demonstrating that phosphorylation of Thr552 markedly reduces PIP_2_ binding. Whereas previous models emphasized disruption or rearrangement of CaM interactions^11,28^, our data support a mechanism in which CaM association is largely preserved while PIP_2_ binding is substantially reduced. Together, these findings suggest that modulation of PIP_2_ interactions may represent a conserved consequence of phosphorylation within the distal B-helix regulatory region of Kv7 channels.

Collectively, our findings support a model in which phosphorylation of Thr552 suppresses Kv7.4 activity primarily through disruption of PIP_2_-dependent regulation rather than by altering channel trafficking or causing dissociation of CaM. Instead, phosphorylation appears to reorganize the local interaction network surrounding the distal B helix polybasic KRK region, reducing PIP_2_ interactions while preserving overall CaM association. These changes shift channel activation toward more depolarized potentials and provide a mechanistic explanation for the observed loss-of-function phenotype. Together, these results identify Thr552 as a critical regulatory node linking phosphorylation-dependent signaling to Kv7.4 gating through modulation of PIP_2_ interactions.

## EXPERIMENTAL PROCEDURES

### Plasmids

The plasmids pcDNA3.1(+)-KCNQ4 (UniProt: P56696), pECFP-C1-CaM, and (pETGQ.H-CaM) were generously provided by Dr. Mark S. Shapiro and pEGFP-C1 was gifted by Dr. James D. Stockand. Sequence analysis revealed an unintended S485F substitution in the KCNQ4 cDNA that was corrected by Q5 site-directed mutagenesis (New England Biolabs). The T552A (phospho-null) and T552D (phosphomimetic) KCNQ4 mutants that were used for electrophysiological experiments were generated from the corrected KCNQ4 construct also using the Q5 site-directed mutagenesis kit. For the co-IP and surface expression studies, N-terminal c-Myc-tagged versions of WT, T552A and T552D KCNQ4 constructs were created by insertion of a c-Myc epitope immediately downstream of the initiating methionine using Q5 site-directed mutagenesis. pEYFP-N1-KCNQ4 constructs were generated using the In-Fusion Cloning kit (Takara Bio Inc, USA). All plasmid sequences were verified by whole plasmid sequencing (Plasmidsaurus, USA).

### Cell culture and transfection

HEK293T cells were procured from the American Type Culture Collection (ATCC) and cultured in Dulbecco’s Modified Eagle’s Medium (Corning; MT10013CV) supplemented with 10% FBS and maintained at 37°C; 5% CO2 in a Heracell VIOS 160i 5% CO2 humidified incubator (ThermoScientific). For electrophysiology recordings, ∼0.5 x 10^6^ cells (< passage number 25) were seeded into 35 mm polystyrene dishes and transfected 24 hours later with 1 µg KCNQ4 plasmid, 0.5 µg pEGFP (as a fluorescent reporter), and 1 µg CFP-CaM for experiments calling for CaM co-transfection, using the Fugene-HD transfection reagent (Promega) according to the manufacturer’s protocol. At around 36 hours post-transfection, transfected cells were trypsinized, and diluted 1/12 into new 35 mm polystyrene dishes, and whole cell recordings conducted 8-24 hours later.

### Peptides, protein constructs, chemicals, and reagents

Peptides were synthesized to 95% purity (Peptide 2.0). The lyophilized peptides were reconstituted in HBS buffer, which consists of 20 mM HEPES and 150 mM NaCl at pH 7.4. To produce untagged calmodulin protein, pETGQ.H-CaM was transformed into BL21 competent cells and induced with 0.5 mM IPTG at 37 °C for 6 h in LB (for binding assays) or minimal essential medium containing [^15^N]-ammonium chloride in D_2_O, (for NMR experiments). CaM was purified using phenyl-Sepharose matrix (Cytiva). The eluent was further purified through a Superdex 75 column on an AKTA FPLC system (GE Healthcare). For apoCaM studies, the protein was exchanged to HBS buffer that had been soaked overnight with Chelex 100 reagent (Bio-Rad) followed by vacuum filtration through a 0.2 uM filter. We refer to this chelexed buffer as “ChHBS.” Glassware and other containers used for apoCaM measurements were pre-rinsed with 10 mM EGTA, followed by 2 x ChHBS rinses, prior to the addition of proteins. The peptides used for apoCaM interactions were reconstituted in ChHBS. The concentrations of the peptides and CaM were determined by SDS-PAGE densitometry compared to known standards of CaM or measuring the absorbance at 280 nm using extinction coefficient 2,980 M^-1^cm^-1^. The PIP_2_ headgroup analogs, PHG-Cy5 and PHG-FL, were purchased from Echelon Biosciences Inc. (Salt Lake City, UT) and dissolved in water at a concentration of 250-311 µM which was diluted to the working concentrations in either HBS or ChHBS buffer.

### Protein Structure Analysis

The protein structures were downloaded from the Protein Data Bank (PDB). Molecular graphics and analyses performed with UCSF Chimera, developed by the Resource for Biocomputing, Visualization, and Informatics at the University of California, San Francisco, with support from NIH P41-GM103311^31^.

### Patch Clamp

Patch clamp experiments were carried out according to the standard protocols outlined elsewhere^21,32^. Bath solutions contained 145 mM NaCl, 4 mM KCl, 1.8 mM CaCl_2_, 0.5 mM MgCl_2_, 10 mM HEPES, and 5 mM D-glucose (pH 7.4 adjusted with NaOH; Osm: 315-325), while the pipette solution contained 140 mM KCl, 1 mM MgCl_2_ and 10 mM HEPES (pH 7.4 adjusted with KOH; Osm: 300-310). For elevated intracellular Ca^2+^ recordings, 50 µM CaCl_2_ was added to the pipette solution without EGTA. For low Ca^2+^ recordings, 10 mM EGTA was added to the pipette solution to buffer intracellular Ca^2+^. The resulting free Ca^2+^ concentration was estimated to be 88 nM using the Ca-EGTA NIST v1.2 calculator (UC Davis Health)^33^. Whole cell patch clamp recording experiments were carried out at room temperature using pulled borosilicate glass pipettes (World Precision Instrument; 1B150F4; resistance of 3-5 megaohms) on isolated GFP-positive cells using an Axopatch™ 200B amplifier (Molecular Devices), digitized using the Digidata 1322A (Molecular Devices), and acquired using the pCLAMP 10 software (Molecular Devices). Recording data was collected at a 10 kHz sample rate and filtered at 5 kHz with a low-pass filter. Series resistance was typically ∼5-10 MΩ and compensated by 80%. Current-voltage (I-V) relationships were obtained from cells held at −80 mV using 2 sec voltage steps from −80 mV to +40 mV in 10-mV increments, followed by a 1 sec tail pulse to −50 mV. Current densities were calculated from currents measured at 1500 ms during each voltage step and normalized to membrane capacitance (pA/pF). The peak current density (I_max_) was determined as the current density at +40 mV for all three variants. To determine the voltage-dependent activation, V_0.5,act_ and the slope factor (*κ*) were obtained by fitting normalized tail current amplitudes with the Boltzmann equation:

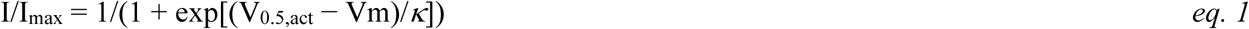

The collected patch-clamp recording data were analyzed using the Clampfit 10.6 software (Molecular Devices, USA), Microsoft Excel (Microsoft), and GraphPad Prism 10 software (Graph Pad). All data was shown as mean ± SD. Statistical analyses were performed in GraphPad Prism 10 using one-way ANOVA followed by Bonferroni’s multiple-comparisons test.

### Cell-surface biotinylation assay

To quantify the surface expression of KCNQ4 channels, 1.5×10^6^ HEK293T cells were seeded into 60mm dishes and transfected using the Fugene-HD transfection reagent (Promega; HD-1000) with 1.5µg of myc tagged-hKCNQ4 and 1.5µg CFP-fused hCaM plasmids, 24 hours post-seeding. The exogenous CFP-tagged CaM (CFP-CaM) was essential in increasing the amount of detectable CaM in HEK293T cells^8^. Cell surface biotinylation and NeutrAvidin pull-down was performed 48 hours post-transfection using a modified protocol adapted from the Pierce™ Cell Surface Protein Biotinylation and Isolation Kit (Thermo Fisher Scientific; 89881). Briefly, cells were incubated with ∼0.5mg/ml of ice cold Sulfo-NHSSS-Biotin for 30 minutes at 4◦C. The biotinylation reaction was then quenched with a buffer containing 100mM glycine and 50Mm Tris HCL, pH 7.4 for 5 minutes at 4◦C, washed twice with ice cold Tris-buffered saline (pH 7.4) and cells lysed for one hour with end-over-end rotation at 4◦C in a lysis buffer containing 50mM Tris pH8.0, 150 mM NaCl, 1% NP40, 5% glycerol, 200 Μm sodium orthovanadate, protease inhibitor cocktail (Thermo Fisher Scientific; 78430), phosphatase inhibitor (Thermo Fisher Scientific; 78420), and either 2mM CaCl2 or 10mM EGTA. The concentration of calcium or EGTA was matched to that used in our patch-clamp recordings. NeutrAvidin was then incubated with 450ul of the cell lysate overnight with end-over-end rotation at 4◦C, followed by three washing steps with ice cold lysis buffer, each lasting 5 minutes, with end-over-end rotation at 4◦C. Elution of biotinylated proteins from the biotin-NeutrAvidin complexes was either done using the 100ul of 5X loading dye (10%SDS, 500mM DTT, 50% Glycerol, 500mM Tris-HCL and 0.5% bromophenol blue dye), or 100ul of the proprietary elution buffer from the surface biotinylation kit. Western blotting was then performed using 1.5mm homemade gels loaded with equal volumes (20 µl) of the biotinylated eluted samples. Green Fluorescent Protein was used as the cytosolic marker. Proteins were transferred to 0.2 µm nitrocellulose membrane using the Trans-Blot® Turbo™ Transfer System (Biorad; USA), incubated with the respective primary and secondary antibodies, and imaged using the Bio-Rad ChemiDoc Imaging System. Primary antibodies: Anti-c-Myc Tag Antibody (Sigma; 06-549-25UG, 1: 1,000), Anti-Green Fluorescent Protein Antibody (Millipore Sigma; MAB3580, 1:1,000). Secondary antibodies: IRDye® 680RD Goat Anti-mouse IgG (Li-COR, Lincoln NE, USA; 926-68071; 1:10,000) and IRDye 800 CW goat anti rabbit (Li-COR; 925-32211; 1:10,000). Band intensities of western blot images from both the lysate and elutions were quantified using ImageJ (NIH, USA).

### Total Internal Reflection Fluorescence Microscopy

To assess whether the T552A and T552D mutations altered surface expression of Kv7.4 channels, Total Internal Reflection Fluorescence (TIRF) microscopy was performed. HEK293T cells were transfected with 0.25 μg of YFP-KCNQ4 plasmids (WT, T552A, or T552D) using Fugene-HD transfection reagent (Promega; HD-1000) in 35-mm glass-bottom dishes (MatTek; P35G-1.5-14-C). Twenty-four hours after transfection, cells were washed three times with Dulbecco’s PBS (ATCC; 30-2200) and maintained in fresh PBS during imaging. All imaging experiments were completed within 20 min of buffer exchange.

Images were acquired using an Olympus IX81 microscope equipped with an Andor iXon Ultra 888 EMCCD camera and a 100× objective. Imaging parameters were held constant for all experiments (laser power, 10 mW; EM gain, 15; exposure time, 0.1 s). All analyses were performed using single-frame images rather than averaged image projections. The number of cells analyzed for each condition is indicated in the corresponding figure legend.

FluoSpheres Carboxylate-Modified Microspheres (Invitrogen; F10720) were used as fluorescence calibration standards. Images were analyzed using ImageJ (NIH). Background fluorescence was determined from nonfluorescent regions of each image and subtracted from both bead and cellular fluorescence measurements. Individual membrane-associated fluorescent cells were manually selected, while out-of-focus or overlapping cells were excluded from analysis. Mean fluorescence intensity was measured for each cell and normalized to the fluorescence intensity of calibration beads acquired during the same imaging session. Relative fluorescence values were pooled for each construct and reported as mean ± SD.

Statistical analyses and graph generation were performed using GraphPad Prism. Differences among groups were evaluated using ordinary one-way ANOVA followed by Tukey’s multiple-comparison test.

### Circular dichroism

Circular dichroism (CD) spectra were collected under two solution conditions. For aqueous measurements, Q4Bx, Q4BxpT, Q4BxAAA, and Q4S6JxA peptides were dissolved to 30 µM in 5 mM KH_2_PO₄ and 5 mM NaCl (pH 7.4) and analyzed in a 0.1-cm quartz cuvette (iCuvettes, QL2HR01). For helix-promoting conditions, Q4Bx, Q4BxpT, and Q4BxAAA were dissolved in the same buffer supplemented with 30% (v/v) isopropanol (IPA).

CD spectra were acquired at room temperature using a J-1500 spectropolarimeter (Jasco, Tokyo, Japan) from 250 to 190 nm with a step size of 0.1 nm. Twenty scans were accumulated for each replicate, and three independent replicates were collected for each peptide. Buffer spectra collected under identical conditions were subtracted to generate baseline-corrected spectra. Secondary structure content was estimated using the ChiraKit analysis platform and SECSA algorithm^22^. Alpha-helical, beta-sheet, and random-coil content were calculated from the baseline-corrected spectra using peptide concentration, path length, and molecular-weight information provided to the software.

### Isothermal Titration Calorimetry

ITC titrations were performed at 20°C using MicroCal VP-ITC microcalorimeter (Malvern Instruments). Titrations were conducted in HBS buffer supplemented with 500 µM CaCl_2_ in both the syringe and cell. Samples were degassed for 30 min. 5 µM CaM was placed in the ITC cell and 35-60 µM peptide was added to the titration syringe. Each ITC experiment consisted of at least 24 injections of 10 µL of titrant, preceded by one 4-µL injection. Data were analyzed with MicroCal Origin version 7.0 using the built-in curve-fitting models. Statistical significance was determined by a two-tailed t test.

### HSQC-NMR

NMR experiments were performed using HSQC (non TROSY) experiments. All experiments were conducted in ChHBS, 1 mM EGTA, and 10% (v/v) D_2_O at 298K on a Bruker Avance 700 NMR spectrometer. The NMR spectrum for 125 µM deuterated ^15^N-apoCaM was recorded then combined with unlabeled Q4B or Q4BphT peptide at 1:1.2 (CaM:peptide) molar ratio in ChHBS or added 2 mM CaCl_2_. The methods were performed similarly to those described elsewhere^34,35^.

The assignments for mammalian apoCaM were shared with us by John Putkey (UT Health, Houston, TX), and the assignments for Ca^2+^/CaM kindly provided by Walter Chazin (Vanderbilt University, Nashville, TN) and Adriaan Bax (NIH, Bethesdsa, MD). The raw spectrometer format data were processed using nmrPipe and nmrDraw in NMRbox^36^. The peaks were calculated and visualized using the POKY software suite and in some cases, the overlays were formatted using Adobe Illustrator^37^.

### Model Building and Molecular Dynamics Simulations

The crystal structure of Apo-Calmodulin (CaM) bound to KCNQ4 helices A and B (Q4AB) was obtained from PDB: 6B8L^9^. The Apo-CaM:Q4AB phosphorylated Thr552 was modeled using CHARMM-GUI^38–41^. To generate Ca^2+^/CaM models, 0.012M CaCl_2_ was added (to achieve a ratio of 1 CaM to 4 Ca^2+^). We generated four models for simulations in total: Apo-CaM:Q4AB unphosphorylated, Apo-CaM:Q4AB phosphorylated, Ca^2+^/CaM:Q4AB unphosphorylated, and Ca^2+^/CaM:Q4AB phosphorylated^38–41^. All phosphorylated and unphosphorylated models were solvated in a TIP3P water box extending 10 Å from the protein in all directions in a periodic boundary conditions^42^. To each model 0.15 M NaCl was added to maintain ionic strength comparable to experimental conditions^43^. All systems were minimized using the steepest descent algorithm, followed by NVT equilibration at 300K temperature using a Nosé-Hoover thermostat and NPT equilibration at 1 bar pressure using Parrinello-Rahman barostat in GROMACS with CHARMM36m forcefields^44,45^. GROMACS was used to perform all-atoms molecular dynamics simulation of all four systems in triplicates using established potential^44^. Simulations used leap-frog integration with 2 fs time-step for 500,000,000 steps, totaling 1 µs per trajectory. Each models were simulated three times, totaling 12 simulations.

### Molecular Dynamics Simulations Analysis

To ensure each simulation has reached structural equilibrium, we measured the Root Mean Square Deviation (RMSD) over time. Once the trajectory has reached a plateau in terms of RMSD value, the simulation is considered to have reached a structural equilibrium. The RMSD values for each simulation was calculated using gmx rms package in GROMACS^46^. The side-chain orientation of Thr552 was characterized by calculating the dihedral angle defined by the atoms C–Cα–Cβ–Oγ1. This analysis was used to identify the occupied rotameric states of Thr552 across all four systems. These dihedral angles were measured in VMD^47^. Histograms of the dihedral angles of Thr552 were fit with a three Gaussian function,

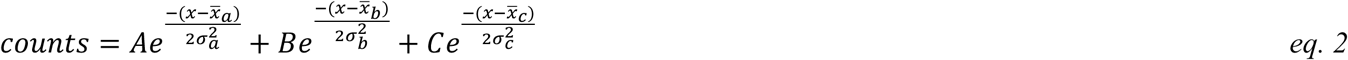

where, A, B, and C are the heights of the Gaussian distributions, *x̅_a_*, *x̅_b_*, and *x̅_c_* are the mean values of each Gaussian distribution, and σ_a_, σ_b_, and σ_c_ are the standard deviations of each Gaussian distribution.

Hydrogen bonds and their occupancies throughout the simulation were identified using VMD plugin with a distance cutoff of 4 Å and an angle cutoff of 80°^47^.

To quantify conformational displacement between CaM and Q4AB, two reference vectors were defined: one connecting Cα atoms of Ala104 and Lys31 of CaM, and another connecting Cα atoms of Leu523 and Leu553 of Q4AB. The former provides a vector along the horizontal axis of CaM as well as CaM-Q4AB complex and the latter provides a vertical axis along B helix as well as CaM-Q4AB complex (**Figure 7K**). The angle between these vectors was calculated to monitor conformational shifts in CaM–Q4AB interactions over the trajectory. Generation of vectors and angle calculations were performed in VMD^47^. An awk script was used to generate mean and standard deviations of the angle between these two vectors.

Backbone dihedral angles φ and ψ for each amino acid were calculated throughout the trajectories. The φ angle for residue n was defined by the atoms C(n−1)–N(n)–Cα(n)–C(n), and the ψ angle was defined by N(n)– Cα(n)–C(n)–N(n+1). The φ and ψ values for each frame of the simulations were plotted on the x- and y-axes, respectively, to generate Ramachandran plots for each amino acid. The presence of α-helices, β-sheets, and left-handed helices was determined based on the population of dihedral angles on the Ramachandran plot. Dihedral angles were measured in Visual Molecular Dynamics (VMD) software^47^.

### Microscale thermophoresis

To create fluorescently labeled CaM (Red-CaM), purified CaM in was conjugated with Red-NHS dye (2^nd^ Generation protein-labeling kit, NanoTemper Technologies, Munich, Germany) in ChHBS and eluted at a concentration of 6.95 µM. For PIP_2_ binding experiments, the fluorescent PIP_2_ analog PHG-Cy5 was purchased (Echelon Biosciences) was dissolved in ChHBS to a stock concentration 250 µM. The peptides were serially diluted 1:2 in ChHBS + 0.5% Tween-20, pH 7.4. ApoCaM binding assays were supplemented with 0.1 mM EGTA with the highest concentration 500 µM. Red-CaM or PHG-Cy5 was added to each dilution to a final concentration of 50 nM. Samples were centrifuged at 500 x *g* for 2 min before loading into standard or premium capillaries (NanoTemper Technologies, MO-K033 or MO-K025). All MST experiments were performed using low-retention tips (Rainin), low-binding microcentrifuge tubes (Eppendorf, Hamburg, G), and non-binding 384-well black plates (Greiner Bio-One) to minimize sample loss and adsorption. Measurements were acquired on a Monolith X (NanoTemper Technologies) using the autodetect settings for excitation power, and medium setting for MST power. Three independent experiments were performed for each interaction (n=3) MST data were fit using the T-jump response, and, and the dissociation constants (K_d_) were determined using PALMIST and visualized using GUSSI software^48–50^. Control experiments are shown in **Figure S5**.

### Co-immunoprecipitation

HEK293T cells transiently expressing myc-KCNQ4 and CFP-CaM were harvested 48 h after transfection and lysed for 30 min in lysis buffer at 4°C. Cell lysates were incubated with 30 µl of Pierce anti-c-Myc magnetic bead slurry for 4 h at 4°C with end-over-end rotation. The beads were washed twice with 700 µl lysis buffer, and bound proteins were eluted using Pierce HA-tag IP/Co-IP elution buffer supplemented with DTT. Eluates were immediately neutralized with 1.0 M Tris-HCl (pH 8.8) and analyzed by SDS-PAGE followed by western blotting. KCNQ4 and CaM were detected using anti-Myc (1:1,000) and anti-CaM (1:1,000) primary antibodies, respectively. IRDye 800CW goat anti-rabbit IgG (1:10,000) was used as the secondary antibody. Band intensities were quantified using ImageJ (NIH). Three independent biological replicates were performed, and CaM intensity was normalized to immunoprecipitated Kv7.4 prior to statistical analysis.

### Fluorescence polarization

Peptides were combined with CaM at equimolar ratios of 200 µM each protein, in HBS buffer supplemented with 500 µM CaCl_2_ or 500 µM EGTA. The protein complexes were serially diluted 1:2, then combined with an equal volume of 200 nM PHG-fluorescein, to achieve a final concentration of 100 nM. The highest final concentration of protein complex was 100 µM protein. The fluorescence was measured at 485/520 nm, with a gain of 60, using a BioTek Synergy microplate reader. Data were fit in GraphPad Prism using the equation:

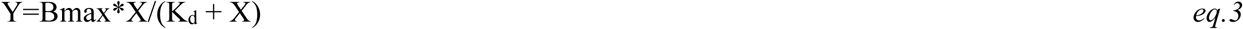

Statistical significance was determined by ordinary 2-way ANOVA with row factor defined as the concentration of the protein complex (combined in equimolar ratios) and column factor defined by the proteins combined in the complex, followed by Tukey’s multiple comparisons t-test. Significant values are indicated as ***, p<0.0001.

### Statistical Analysis

All data are presented as mean ± SD unless otherwise indicated. Statistical analyses were performed using GraphPad Prism 10 or GraphPad Prism 11. Specific statistical tests used for each experiment are described in the corresponding methods section and figure legends. A *p* value <0.05 was considered statistically significant.

## Data Availability

The data supporting the findings of this study are contained within the contents of this article. The datasets generated during this study will be freely provided by the corresponding author upon request.

This article contains supporting information.

## Acknowledgments

We thank James D. Stockand, Ph.D. for valuable guidance and resources during the early stages of this project and Bin Dong for access to TIRF microscopy instrumentation. We thank Chad Brautigam, Ph.D. for technical expertise and guidance in biophysical analyses. We are grateful to Kristin Cano and Akash Bhattacharya for assistance and expertise in NMR experiments. We also thank Cynthia Veliz, Ava Marie McCargo, Jaimeen Patel, Emily Walsh, and Elric Pott for technical assistance and data collection. Finally, we thank Mark S. Shapiro, Ph.D. and Carlos Castaneda, PhD for valuable discussions and feedback on this work.

## Author contributions (CRediT)

Conceptualization: C.R.A.

Methodology: C.R.A., P.P., D.G., A.B., C.O.

Investigation: P.P., C.O., W.M., D.S., C.R.A.

Formal Analysis: P.P, W.M. CRA

Writing – Original Draft: P.P., W.M. C.R.A

Writing – Review & Editing: C.R.A., P.P., D.G., C.O.

Supervision: C.R.A., D.G.

Software: C.R.A., D.G

Project Administration: C.R.A.

Funding Acquisition: C.R.A., DG

