## Supplemental Information for "Phosphorylation of the Kv7.4 B helix reorganizes calmodulin interactions and reduces PIP_2_ binding"

Sequence Name: Q4Bx

|  |  |
| --- | --- |
|  | 004444999CEEEEDDBBCA7666663333000 |
| 524 | -----+-----+-----+-----+ |
| Residue | DDIMPAVKTVIRSIRILKFLVAKRKFKETLRPY |
| 1-10 | 1112333333332222221111 |
| 1-5-10 | 1112222222221111111 |
| BASIC_1-5-10 |  |
| 1-12 | 11111112222111122211111111 |
| 1-14 | 11112222222222222111111111 |
| 1-8-14 | 11112222222221111 |
| 1-5-8-14 | 11112222222221111 |
| BASIC_1-8-14 |  |
| 1-16 | 1111111111111111 |
| IQ |  |
| IQ-LIKE |  |
| IQ-2A |  |
| IQ-2B |  |
| IQ-UNCONVENTIONAL |  |

**Supplemental Figure S1. CaM binding motif prediction of the Kv7.4 B helix.** The sequence of the B helix was entered into the Calmodulin database and meta-analysis predictor (https://cam.umassmed.edu/) to predict the residues that interact with CaM.

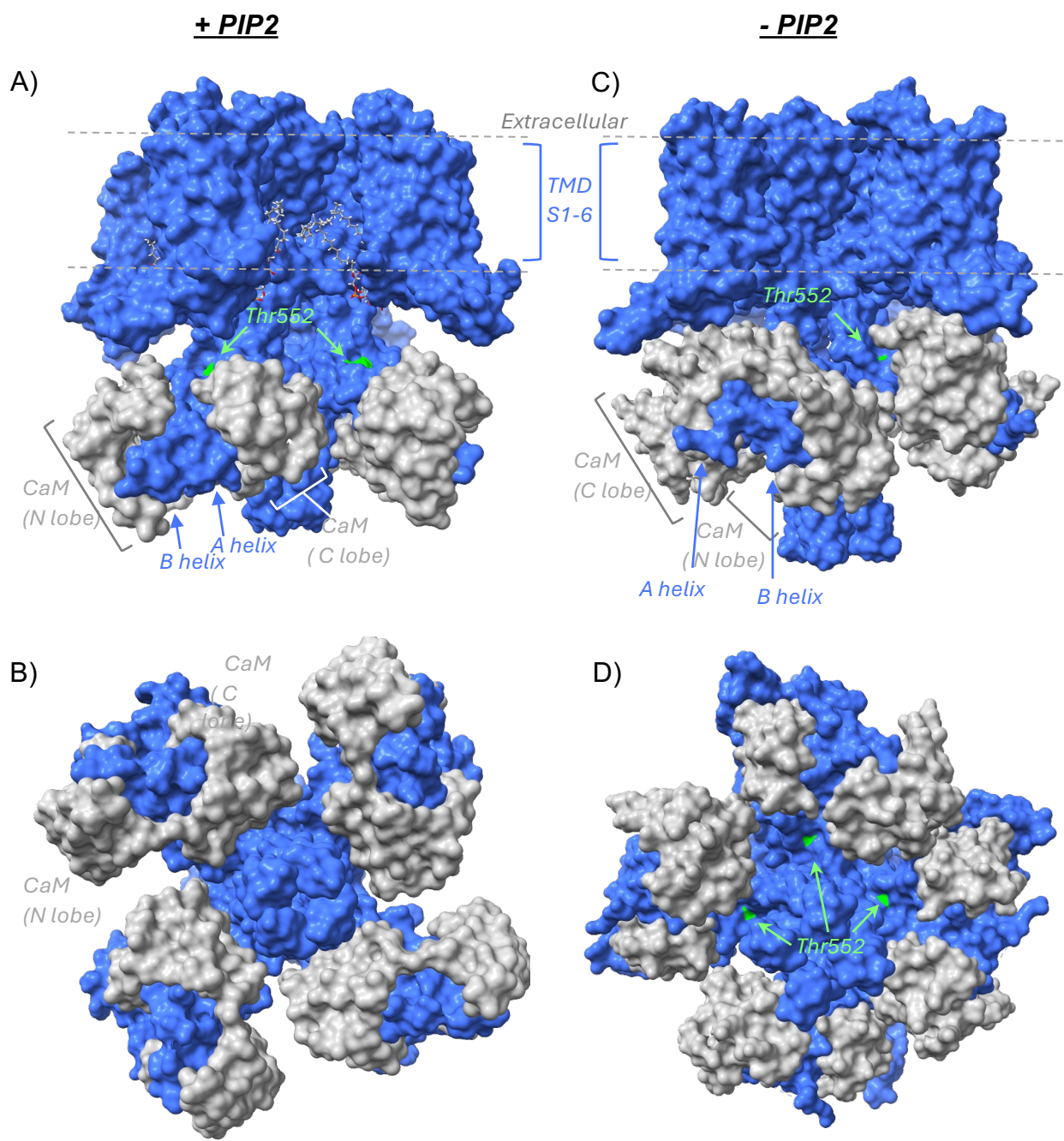

**Supplemental Figure S2.** Comparison of the surface exposure of Thr552 in the CaM-KCNQ4 complex in the PIP2-bound (PDB: 7VNP, *left*) and PIP2-free (PDB: 7VNQ, *right*) states. Top panels show the side view and bottom panels show the view from the bottom, intracellular side of the channels. The structures were downloaded from the Protein Databank and rendered in Chimera. Each of the 4 KCNQ4 subunits is colored blue and CaM is colored grey. Thr552 is colored green. The PIP2 molecules are modeled as stick style and dotted line indicates the channel transmembrane domains.

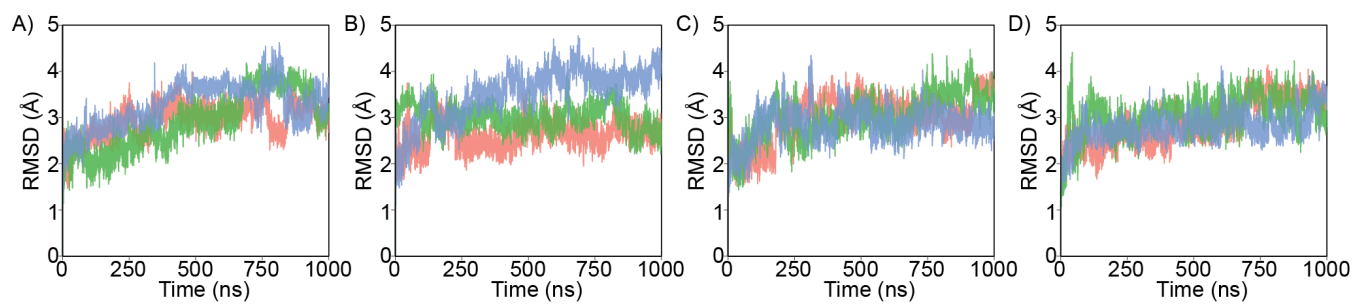

**Supplemental Figure S3. Root Mean Square Deviations of all Systems.** Root mean square deviations of all Cα atoms of the (A) ApoCaM:Q4AB unphosphorylated, (B) ApoCaM:Q4AB phosphorylated, (C) Ca<sup>2+</sup>/CaM:Q4AB unphosphorylated, and (D) Ca<sup>2+</sup>/CaM:Q4AB phosphorylated systems in triplicates.

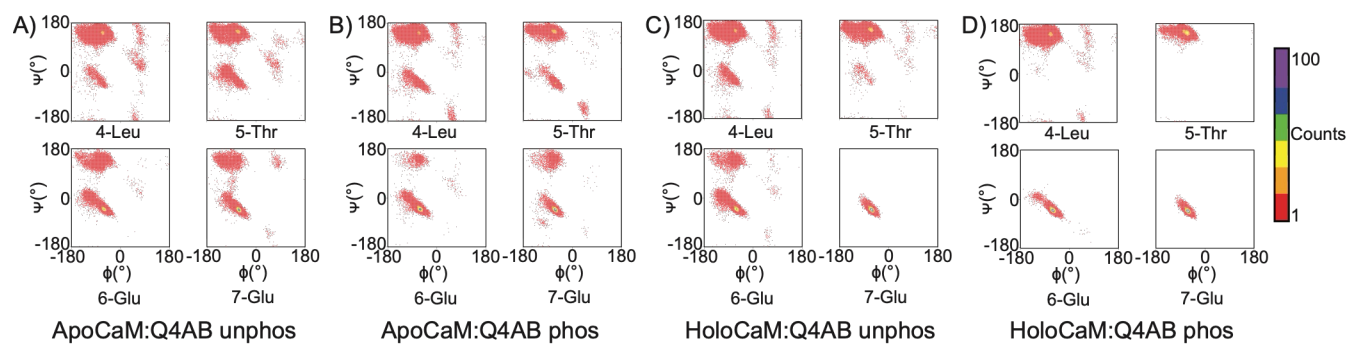

**Supplemental Figure S4. Changes in backbone conformational preferences of N-terminal residues of CaM.** Changes in backbone conformational preferences (φ-ψ distributions) of residues 5–8 of CaM in (i) ApoCaM:Q4AB unphosphorylated system, (ii) ApoCaM:Q4AB phosphorylated system, (iii) $\text{Ca}^{2+}$/CaM:Q4AB unphosphorylated system, (iv) $\text{Ca}^{2+}$/CaM:Q4AB phosphorylated system.

### Supplemental data:

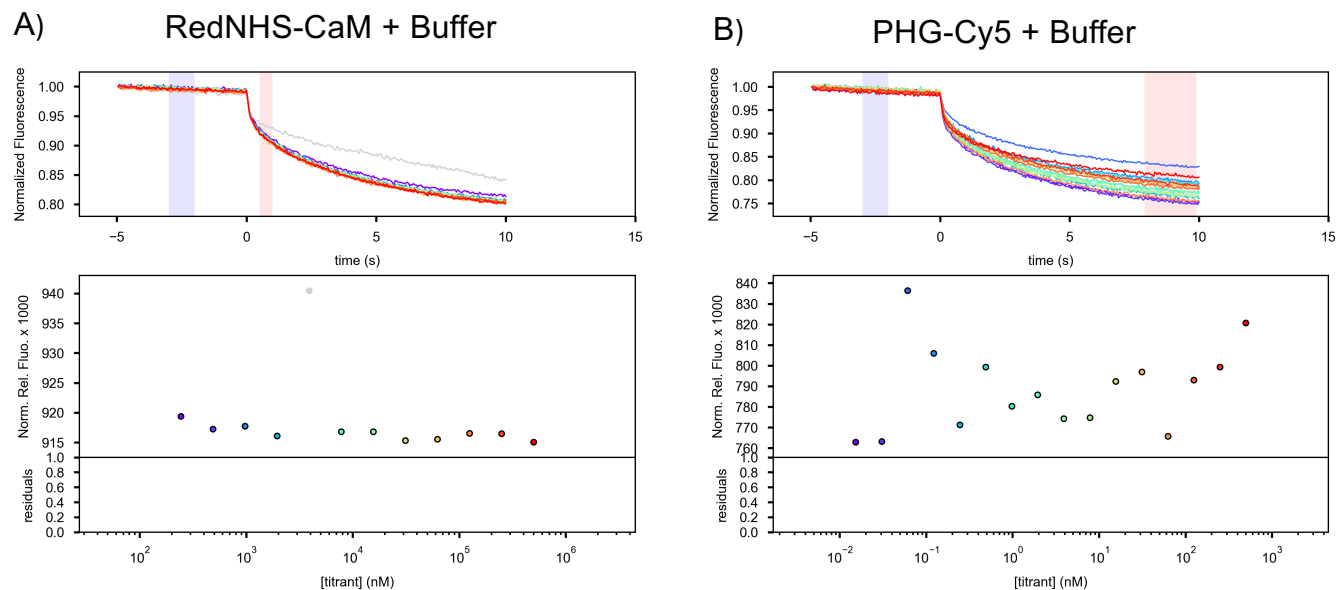

**Supplemental Figure S5. Control MST experiments for apoCaM binding Q4 peptides. (A)** Control experiments for RedNHS-CaM titrated with Ca<sup>2+</sup>-free buffer (chHBS + 0.1 mM EGTA, 0.05% Tween 20, pH = 7.4), showing no binding was detected. **(B)** Control experiments for PHG-Cy5 titrated with Ca<sup>2+</sup>-free buffer (ChHBS+0.1 mM EGTA, 0.05% Tween 20, pH = 7.4), showing no binding was detected.

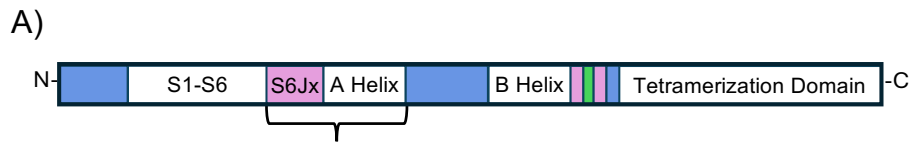

**Q4A peptide:** 336 -----EKRRMPAANLIQAAWRLYSTD
**Q4S6JxA peptide:** 328 **EQHRQKHF**EKRRMPAANLIQAAWRLYSTD

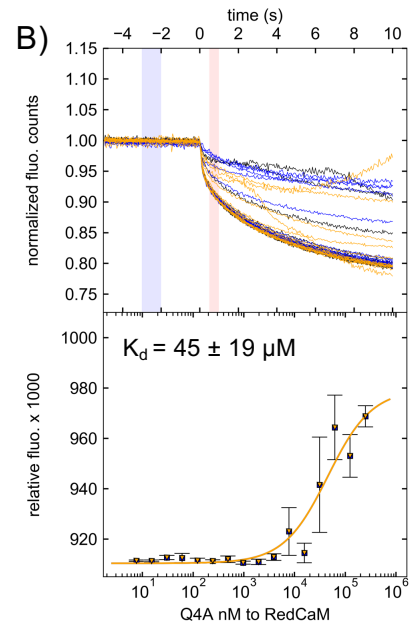

**Supplemental Figure S6. Binding affinities between apoCaM and Q4A short peptide.** Peptide sequence showing the differences in amino acid sequences between the Q4A peptide and the Q4S6JxA peptide (A). MST fluorescence graphs and thermograms for the Q4A peptide titrated into 50 nM of Red-NHS-apoCaM (20 mM HEPES, 150 mM NaCl, 0.1 mM EGTA, 0.05% Tween 20, pH = 7.4), analyzed at 650 nm using the T-jump analysis mode in PALMIST. The data was collected with $n=3$ in the MST low-binding premium capillary. Results are shown as Mean ± SD.

| Table ST1. Channel activity results. |  |  |  |  |  |
| --- | --- | --- | --- | --- | --- |
|  |  | Activation (tail currents) |  | Peak current |  |
| Relative Ca<sup>2+</sup> levels | Kv7 construct | V<sub>0.5 act</sub> [mV] | k<sub>act</sub> [mV] | I<sub>max</sub> at +40 mV [pA/pF] | n |
| without CaM overexpression (-CaM) |  |  |  |  |  |
| 50 uM Ca<sup>2+</sup> | Q4-WT | -24.60 ± 6.282 | 12.36 ± 1.731 | 96.40 ± 33.13 | 8 |
|  | Q4-T552A | -22.75 ± 5.799 | 11.29 ± 0.9821 | 56.90 ± 22.87** | 9 |
|  | Q4-T552D | -13.74 ± 9.498* | 8.019 ± 1.410*** | 22.65 ± 9.114**** | 7 |
| 10 mM EGTA | Q4-WT | -25.67 ± 2.5 | 9.48 ± 1.19 | 247.4 ± 131.9 | 8 |
|  | Q4-T552A | -27.74 ± 2.69 | 9.90 ± 1.65 | 238.3 ± 117.9 | 10 |
|  | Q4-T552D | -4.91 ± 9.21**** | 8.43 ± 1.31 | 64.10 ± 36.73 ** | 11 |
| with CaM overexpression (+CaM) |  |  |  |  |  |
| 50 uM Ca<sup>2+</sup> | Q4-WT | -29.95 ± 7.50 | 11.70 ± 2.70 | 96.13 ± 42.68 | 12 |
|  | Q4-T552A | -31.95 ± 7.92 | 10.68 ± 1.70 | 101.8 ± 41.46 | 8 |
|  | Q4-T552D | -10.99 ± 3.50*** | 14.30 ± 4.14 | 34.34 ± 17.40** | 7 |
| 10 mM EGTA | Q4-WT | -31.58 ± 5.33 | 9.94 ± 2.24 | 393.8 ± 183.7 | 8 |
|  | Q4-T552A | -29.35 ± 6.20 | 9.59 ± 3.17 | 408.2 ± 115.2 | 9 |
|  | Q4-T552D | -11.11 ± 4.71**** | 9.52 ± 1.56 | 53.52 ± 20.58 **** | 8 |

**Supplemental Table ST1. Electrophysiology results of Kv7.4-WT, phospho-null and phosphomimetic constructs in Ca<sup>2+</sup> and EGTA-buffered conditions in the absence and presence of CaM in HEK293T cells.** Voltage-dependence of activation parameters were obtained from Boltzmann fitting of normalized tail currents activation curves. Peak current density (I<sub>max</sub>) at +40 mV. Data are presented as mean ± SD. Statistical comparisons were performed using one-way ANOVA followed by Bonferroni's multiple comparisons test. n indicates the number of cells analyzed. V<sub>0.5 act</sub>, half-maximal voltage of activation, 'k', slope factor of activation, and I<sub>max</sub>, peak current density.

**Supplemental Table ST2.** Residues with peak height >2SD of the mean free CaM.

| # peaks ≥ 2SD: | 20 | 65 | 126 | 106 |
| --- | --- | --- | --- | --- |
|  | apoCaM vs Q4BpHT | apoCaM + Q4Bext | Ca<sup>2+</sup>/CaM + Q4BpHT | Ca<sup>2+</sup>/CaM + Q4Bext |
|  | 95.D.N-NH | 5.T.N-NH | Q3.N-NH | L4.N-NH |
|  | 100.I.N-NH | 13.K.N-NH | L4.N-NH | T5.N-NH |
|  | 101.S.N-NH | 16.F.N-NH | T5.N-NH | A10.N-NH |
|  | 102.A.N-NH | 17.S.N-NH | E6.N-NH | E11.N-NH |
|  | 108.V.N-NH | 20.D.N-NH | E7.N-NH | F12.N-NH |
|  | 111.N.N-NH | 22.D.N-NH | I9.N-NH | K13.N-NH |
|  | 113.G.N-NH | 24.D.N-NH | A10.N-NH | A15.N-NH |
|  | 128.A.N-NH | 25.G.N-NH | E11.N-NH | S17.N-NH |
|  | 129.D.N-NH | 47.E.N-NH | F12.N-NH | L18.N-NH |
|  | 131.D.N-NH | 50.D.N-NH | K13.N-NH | F19.N-NH |
|  | 134.G.N-NH | 51.M.N-NH | E14.N-NH | D20.N-NH |
|  | 136.V.N-NH | 52.I.N-NH | A15.N-NH | K21.N-NH |
|  | 137.N.N-NH | 65.F.N-NH | F16.N-NH | D22.N-NH |
|  | 138.Y.N-NH | 71.M.N-NH | S17.N-NH | G23.N-NH |
|  | 139.E.N-NH | 76.M.N-NH | L18.N-NH | D24.N-NH |
|  | 140.E.N-NH | 78.D.N-NH | F19.N-NH | G25.N-NH |
|  | 141.F.N-NH | 79.T.N-NH | D20.N-NH | T26.N-NH |
|  | 145.M.N-NH | 80.D.N-NH | K21.N-NH | I27.N-NH |
|  | 147.A.N-NH | 81.S.N-NH | D22.N-NH | T28.N-NH |
|  | 148.K.N-NH | 86.R.N-NH | G23.N-NH | T29.N-NH |
|  |  | 87.E.N-NH | D24.N-NH | K30.N-NH |
|  |  | 88.A.N-NH | G25.N-NH | E31.N-NH |
|  |  | 89.F.N-NH | T26.N-NH | L32.N-NH |
|  |  | 90.R.N-NH | I27.N-NH | G33.N-NH |
|  |  | 91.V.N-NH | T28.N-NH | T34.N-NH |
|  |  | 92.F.N-NH | T29.N-NH | V35.N-NH |
|  |  | 93.D.N-NH | E31.N-NH | M36.N-NH |
|  |  | 95.D.N-NH | L32.N-NH | R37.N-NH |
|  |  | 97.N.N-NH | G33.N-NH | G40.N-NH |
|  |  | 100.I.N-NH | V35.N-NH | Q41.N-NH |
|  |  | 101.S.N-NH | M36.N-NH | T44.N-NH |
|  |  | 102.A.N-NH | R37.N-NH | E47.N-NH |
|  |  | 105.L.N-NH | S38.N-NH | M51.N-NH |
|  |  | 107.H.N-NH | L39.N-NH | I52.N-NH |
|  |  | 109.M.N-NH | G40.N-NH | N53.N-NH |
|  |  | 111.N.N-NH | Q41.N-NH | E54.N-NH |
|  |  | 112.L.N-NH | T44.N-NH | V55.N-NH |
|  |  | 113.G.N-NH | E45.N-NH | D56.N-NH |
|  |  | 114.E.N-NH | A46.N-NH | A57.N-NH |
|  |  | 117.T.N-NH | E47.N-NH | D58.N-NH |
|  |  | 119.E.N-NH | M51.N-NH | G59.N-NH |
|  |  | 121.V.N-NH | I52.N-NH | N60.N-NH |
|  |  | 122.D.N-NH | N53.N-NH | G61.N-NH |
|  |  | 124.M.N-NH | E54.N-NH | T62.N-NH |
|  |  | 125.I.N-NH | V55.N-NH | I63.N-NH |
|  |  | 126.R.N-NH | D56.N-NH | D64.N-NH |
|  |  | 127.E.N-NH | A57.N-NH | F65.N-NH |
|  |  | 128.A.N-NH | D58.N-NH | E67.N-NH |
|  |  | 129.D.N-NH | G59.N-NH | F68.N-NH |
|  |  | 130.I.N-NH | N60.N-NH | T70.N-NH |
|  |  | 131.D.N-NH | G61.N-NH | M71.N-NH |
|  |  | 133.D.N-NH | T62.N-NH | M72.N-NH |
|  |  | 134.G.N-NH | I63.N-NH | A73.N-NH |
|  |  | 135.Q.N-NH | D64.N-NH | R74.N-NH |
|  |  | 136.V.N-NH | F65.N-NH | Q75.N-NH |
|  |  | 137.N.N-NH | F68.N-NH | M76.N-NH |
|  |  | 138.Y.N-NH | L69.N-NH | K77.N-NH |
|  |  | 139.E.N-NH | T70.N-NH | D78.N-NH |
|  |  | 140.E.N-NH | M71.N-NH | T79.N-NH |
|  |  | 141.F.N-NH | M72.N-NH | D80.N-NH |
|  |  | 142.V.N-NH | A73.N-NH | E82.N-NH |
|  |  | 143.Q.N-NH | R74.N-NH | E83.N-NH |
|  |  | 145.M.N-NH | Q75.N-NH | I85.N-NH |
|  |  | 147.A.N-NH | M76.N-NH | R86.N-NH |
|  |  | 148.K.N-NH | K77.N-NH | A88.N-NH |

**Supplemental Table ST2.** Comparison of HSQC-NMR CaM peaks. All peaks listed had a peak height that changed >2SD from the mean of unbound apoCaM, *left columns* or unbound Ca<sup>2+</sup>/CaM, *right columns*. The cells colored orange indicate the changed peaks that were common for apoCaM after titrated with phosphorylated peptide compared to unphosphorylated peptide. Likewise, the cyan colored cells indicate common peaks between the peptides of Ca<sup>2+</sup>/CaM. The numbers at the top of each column indicate the total number of changed peaks for each group.
